# Functional loss of *rffG* and *rfbB,* encoding dTDP-glucose 4,6-dehydratase, changes colony morphology, cell shape, motility and virulence in *Salmonella* Typhimurium

**DOI:** 10.1101/2023.06.27.546680

**Authors:** Subhashish Chakravorty, Pip Banerjee, Joel P. Joseph, Sanmoy Pathak, Taru Verma, Mrinmoy Das, Dipankar Nandi

## Abstract

Lipopolysaccharide (LPS) O-antigen and enterobacterial common antigen (ECA) play crucial roles in maintaining the structural integrity of the outer membrane in Gram-negative bacteria. Previous studies conducted with either LPS or ECA mutants have highlighted the importance of these cell surface polysaccharides in the physiology of *Salmonella enterica* serovar Typhimurium (*S*. Typhimurium). However, the functional consequences resulting from the abrogation of both O-antigen and ECA synthesis in *S*. Typhimurium are not well studied. In the present study, we generated single and double gene-deleted mutants of *rffG* and *rfbB*, which are paralogs, encoding dTDP-glucose 4,6-dehydratase that catalyze steps in the synthesis of both O-antigen and ECA. The functional loss of both *rffG* and *rfbB* (Δ*rffG*Δ*rfbB*), but not in single gene-deleted strains, results in a round cell morphology, smaller colony formation and altered LPS profile. In addition, the Δ*rffG*Δ*rfbB* strain displays defects in outer membrane permeability, causing hypersensitivity to bile and cell wall targeting antibiotics, e.g., meropenem and polymyxin B. Transcriptomic analysis identified flagellar and SPI-1 pathway to be highly down-regulated in the Δ*rffG*Δ*rfbB* strain which leads to impaired swimming and swarming motility and lower adhesion and invasion of HeLa cells. Importantly, the Δ*rffG*Δ*rfbB* strain is less proficient in colonizing Peyer’s patches, spleen and liver, is unable to induce pro-inflammatory cytokines and is attenuated in both the oral and intra-peritoneal models of *S*. Typhimurium infection in mice. Overall, this study highlights the importance of *rffG* and *rfbB* in maintaining cell wall integrity, colony and cellular morphology, motility and virulence in *S*. Typhimurium.

## Introduction

Salmonellosis is a global public health concern, especially with the advent of anti-microbial resistant (AMR) bacterial strains on the rise. The treatment of infections caused by *Salmonella* with the current generation of antibiotics continue to be a challenge owing to the presence of the outer membrane which is impervious to the action of several antibiotics that are otherwise effective against Gram-positive pathogens (1). The outer membrane of Gram-negative bacteria is made up of lipopolysaccharide (LPS) and outer membrane proteins (OMPs). The presence of the LPS molecules provide a permeability barrier which excludes toxic substances such as large hydrophobic antibiotics (2). In *S*. Typhimurium, the LPS molecules that are present on the outer leaflet consists of three structural units, the Lipid A, the core, and O-antigen repeating units which are made up of tetra-saccharide units of D-mannose, L-rhamnose, D-galactose and D-abequose (3). Another antigenic feature which is exclusively present on the outer membrane of species from the order Enterobacterales, is the enterobacterial common antigen (ECA). ECA is made up of repeating trisaccharide units composed of 4-acetamide-4,6 dideoxy D-galactose, N-aceyl-D-mannosaminuronic acid and N-acetyl-D-glucosamine (4). The LPS, along with ECA, provides structural integrity to the outer membrane which is vital for the bacteria as their loss is associated with significant fitness cost (5, 6).

Agents which compromise the integrity of the outer membrane hold the promise to be a potent therapeutic option. However, there is no drug available to efficiently target the components required for the assembly of the outer membrane of bacteria without having toxic effects on the human body (7, 8). Thus, a potential drug target which is very specific to the pathogen without having any adverse side effect on the host is highly desirable. One such potential drug target which has garnered significant interest in recent times, owing to its ubiquitous presence in both Gram-positive and Gram-negative pathogens is the rhamnose biosynthesis pathway. The L-rhamnose pathway is found in both plants and bacteria and is a well-characterized pathway that has been studied extensively over the past three decades. However, in recent times it has gained considerable interest owing to its potential as a therapeutic target. L-rhamnose is a component of bacterial cell wall and capsule of many pathogenic bacteria and the pathway is highly conserved in both Gram-positive and Gram-negative bacteria. L-rhamnose is a common component of the O-antigen of lipopolysaccharide (LPS) of various Gram-negative bacteria like *Escherichia coli*, *Salmonella enterica* and *Shigella flexneri* (9–11). The genes encoding for the O-antigen biosynthesis pathway are present as a gene cluster called the *rfb* region, which is about 20kb in length and comprises of several genes (12). It is well established that pathways involved in O-antigen biosynthesis are crucial for pathogenesis of several Gram-positive and Gram-negative bacterial species (13).

*Salmonella enterica* serovar Typhimurium (*S*. Typhimurium) must overcome a wide array of host-mediated antimicrobial responses to establish successful infection in the host. The outer membrane in these bacteria is an important structural feature which enables it to cope with myriad environmental and antimicrobial stress responses. LPS along with ECA forms essential components of immunodominant antigens for these Gram-negative pathogens. Therefore, studying bacterial mutants deficient in LPS and ECA biosynthesis pathway offers a promising area of research. The genes, *rffG* and *rfbB* encode the enzyme, dTDP-D-glucose 4,6-dehydratase which catalyzes the intermittent steps of both O-antigen and ECA biosynthesis and deletion of both these genes render the pathogen incapable of synthesizing these molecules (14). RfbB (RmlB) functions as a homodimer and possess a Rossmann fold at the N-terminal domain (15). The homodimer with monomer association principally occurs through the hydrophobic interactions via a four-helix bundle (16, 17). Each monomer in the enzyme exhibits an α/β structure that can be divided into two domains. The larger N-terminal domain binds the nucleotide cofactor NAD+ and consists of a seven-stranded β-sheet surrounded by α-helices. The C-terminal domain is responsible for binding the sugar substrate, dTDP-D-glucose and contains four β-strands and six α-helices. The two domains meet to form a cavity in the enzyme. The highly conserved active site, Tyr-X-X-X-Lys catalytic couple and the Gly-X-Gly-X-X-Gly motif at the N terminus characterize RmlB as a member of the short-chain dehydrogenase/reductase (SDR) family. Importantly, mutations in the TDP-Glucose 4,6-Dehydratase (TGDS) encoding gene in humans are known to cause Catel-Manzke syndrome which is characterized by abnormalities in the index finger (18).

Previously, our laboratory in collaboration with Prof Parag Sadhale’s lab in MCBL, IISc, had reported the functional roles of UDP-glucose 4,6 dehydratase in *Candida albicans* (19). Previous studies conducted with the single gene-deleted mutants of either O-antigen or ECA have shown the importance of these two proteins on the physiology of *S*. Typhimurium (20–22). However, no study has been undertaken till date to provide a detailed investigation of the effects of the loss of both O-antigen and ECA in *S*. Typhimurium. With this objective in mind, we initiated this study. To this end, we generated single and double gene-deleted mutants of *rffG* and *rfbB* to study the effect of functional loss of dTDP-glucose 4,6-dehydratase. The broad objective of this study was to understand the consequences of the loss of O-antigen and the ECA and its effects on the physiology and virulence of *S*. Typhimurium.

## Results

### Multiple Sequence Alignment (MSA) of RffG and RfbB protein homologues indicate presence of a conserved YXXXK motif

The genes under investigation, *rffG* and *rfbB* are paralogs, encode the enzyme dTDP-glucose 4,6-dehydratase and are organized as part of distinct operons (Figure S1). In order to obtain homologs, the protein FASTA sequences of RfbB (361aa; locus id: STM14_2591) and RffG (355aa; locus id: STM14_4720) from *Salmonella* Typhimurium 14028s were retrieved from NCBI. These sequences were then independently used as query sequences to perform protein BLAST (using the NCBI BLASTp suite) against the target non-redundant protein sequence database. The search set was curated and limited to records from reference genomes of only selected prokaryotic and eukaryotic representative organisms and the BLAST was performed with default algorithm parameters. The hits obtained were further filtered and only those with highest percentage identity in each reference genome were retrieved for further analysis. FASTA sequences producing significant alignment with the input query sequence were downloaded and used as an input in CLUSTAL Omega to generate alignments (23–25). Subsequently, JalView was used to highlight important features in the alignment, like the conserved residues and to generate a consensus sequence (26).

A motif scan with the RfbB sequence on the ExPasy suite ProSite, indicated that the protein belongs to the SDR superfamily, as indicated earlier (16). Earlier reports also suggested that the SDR family is a diverse group of enzymes with residue identity of 15-30%. The crystal structure of dTDP-D-Glucose 4,6-dehydratase (RfbB or RmlB) from *S.* Typhimurium, the second enzyme in the dTDP-l-rhamnose pathway was first published in 2001 (27). The crystal structure shows that there are two regions in the active site: the cavity created by the NAD+ binding region and the dTDP-D-glucose-binding regions (27). We observed that eukaryotic homologs have a very low degree of sequence identity ranging between 25-40%, possibly suggesting functional divergence (data not shown). However, the one strictly conserved residue in this extended SDR family is the Tyr (residue 167), part of the YXXXK motif and is also the substrate binding site. Thr133 and Lys171 in RfbB are also highly but not absolutely conserved. We observed that Ser often conservatively substitutes the Thr at position 133. Other highly conserved residues include the three glycine residues, with motif (Gly-X-Gly-X-X-Gly) located close to the N terminus (at positions 8, 10 and 13 in RmlB). It is known that these residues are involved in NAD^+^ binding. These conserved residues in the SDR family suggest that despite their different enzymatic activities, the common chemical step, i.e., transfer of hydride, largely dictates the enzyme structure.

While the SDR signature motif (LftettayapSspYSASKASSdHLVrAWR) slightly varied in the RffG sequence, it did retain the active site Tyr and Lys residues (Figure S2a). The other residues that are known to make contacts with the nucleotide sugar are Thr133, Asp134, Glu135, Asn196, Arg231 and Asn266. Notably, amino acid variations were observed in *Shigella, Mycobacterium, Drosophila* and *Saccharomyces* species (Figure S2). Mechanistic roles of the residues Thr133, Tyr167 and Lys171 in the catalytic motif of the enzyme has been studied through mutagenesis and steady-state kinetics analysis in *Escherichia coli* (28). Overall, it was interesting to observe the presence of homologs of this enzyme from *E*. *coli* to humans.

### *S*. Typhimurium Δ*rffG*Δ*rfbB* strain shows higher optical density (OD) and a delayed lag phase compared to the WT and single mutant strains

To better understand the functional roles of *rffG* and *rfbB* in *S*. Typhimurium, single and double gene-deleted mutants were generated using the one step gene deletion strategy (29). To determine whether the deletion of *rffG*, *rfbB* or deletion of both the genes had any effect on the growth of *S*. Typhimurium, growth assays were performed. WT, Δ*rffG*, Δ*rfbB* and Δ*rffG*Δ*rfbB* strains were grown in a nutritionally rich medium (LB) and aliquots were obtained and the OD at 600 nm quantified at various time points. As observed, WT and the single mutants displayed similar growth kinetics, however, the Δ*rffG*Δ*rfbB* strain was found to attain a higher OD when compared to other strains from mid-exponential phase till the stationary phase (Figure S3a). Notably, introducing the wildtype copy of either *rffG* or *rfbB* restored the OD to WT levels (Figure S3b).

To further investigate whether the higher OD displayed by the Δ*rffG*Δ*rfbB* strain corresponded with the actual number of viable cells, appropriate dilutions were plated on agar plates at various time points along the growth curve. Cellular viability between the wild type and the single gene-deleted mutants were not significantly different, however, the Δ*rffG*Δ*rfbB* strain at 3 hours showed lesser CFU when compared to other strains (Figure S3c). After the initial time point of 3 hours, we did not observe any difference in viability among the four strains. This implied that apart from demonstrating a prolonged lag phase in liquid medium, there were no other major growth differences in the Δ*rffG*Δ*rfbB* strain compared to the WT and the single gene-deleted mutants. Complementing with the WT copy of either of the genes restored the growth phenotype of the Δ*rffG*Δ*rfbB* strain (Figure S3d).

### *S*. Typhimurium Δ*rffG*Δ*rfbB* forms smaller colonies and a distinct colony morphology on LB agar plates

Interestingly, we also observed that the colonies formed by Δ*rffG*Δ*rfbB* strain on LB agar plates were considerably smaller in size when compared to the WT or the single gene-deleted mutants (Figure 1a). Complementing with the WT copy of either of the genes restored the colony size of the Δ*rffG*Δ*rfbB* to that of the WT (Figure 1a and Figure 1c). This indicated that the Δ*rffG*Δ*rfbB* strain was more compromised in growth on solid media than in the liquid media. Also, we noted that the colony texture of the Δ*rffG*Δ*rfbB* strain appeared much smoother compared to the WT and the single gene-deleted strains (Figure 1b).

**Figure 1.**
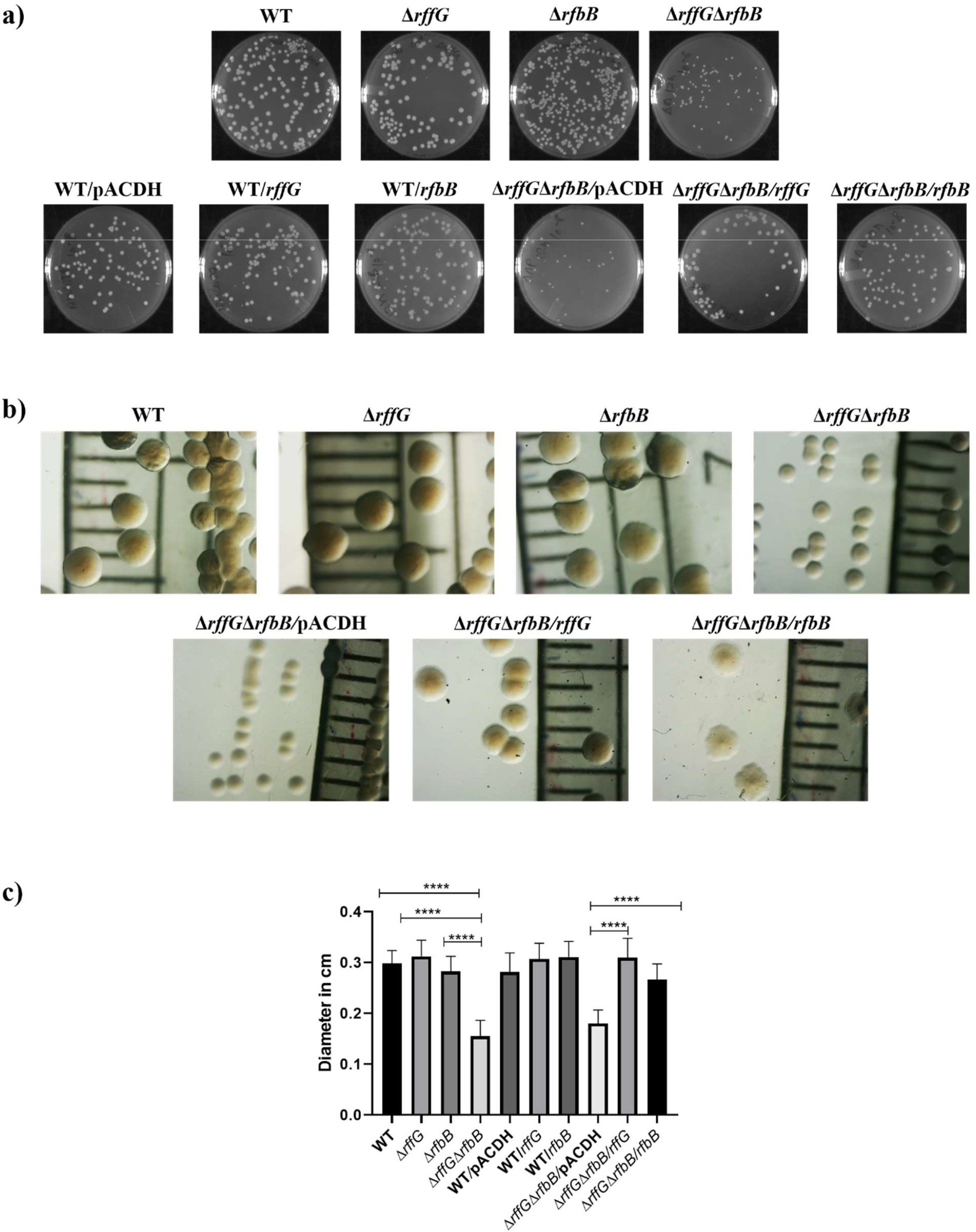
*S*. Typhimurium Δ*rffG*Δ*rfbB* strain forms smaller colonies on LB agar. **(a)** *S*. Typhimurium WT, Δ*rffG*, Δ*rfbB*, Δ*rffG*Δ*rfbB* and complemented strains were plated on LB agar plates, and their images were captured after 16 hours of incubation at 37°C. **(b)** Representative images depicting individual colony morphology of the above-mentioned strains obtained through a light microscope. **(c)** Quantification of the colony size of WT, Δ*rffG*, Δ*rfbB* and Δ*rffG*Δ*rfbB* and the complemented strains on LB agar plates. Statistical analysis was performed using one-way ANOVA, where \*\*\*\**p* < 0.0001. Data are representative of 3 independent experiments plotted as mean ± SEM

### *S*. Typhimurium Δ*rffG*Δ*rfbB* have distinct cell morphology as revealed by Atomic force microscopy (AFM)

Furthermore, we examined whether there were any differences in the cellular morphology among the different strains. To gain morphological insights at a single cell resolution, AFM was performed on the different strains. AFM based techniques are now been routinely used for the multiparametric analysis of cellular surfaces (30). AFM imaging was performed at different time points of bacterial growth i.e., 3, 6 and 12 hours (Figures 2 and S4). The cellular width of the Δ*rffG*Δ*rfbB* strain was found to be greater than the WT strain and single gene-deleted mutants. At the 6^th^ and the 12^th^ hour, most of the cells of the Δ*rffG*Δ*rfbB* strain exhibited a round or a spherical morphology (Figure S4b). Quantification of the cellular width revealed that the Δ*rffG*Δ*rfbB* strain was significantly wider compared to the WT strain. This phenotype was completely restored upon the expression of the WT copy of either *rfbB* or *rffG in trans* (Figure S4a and Figure S4c).

**Figure 2.**
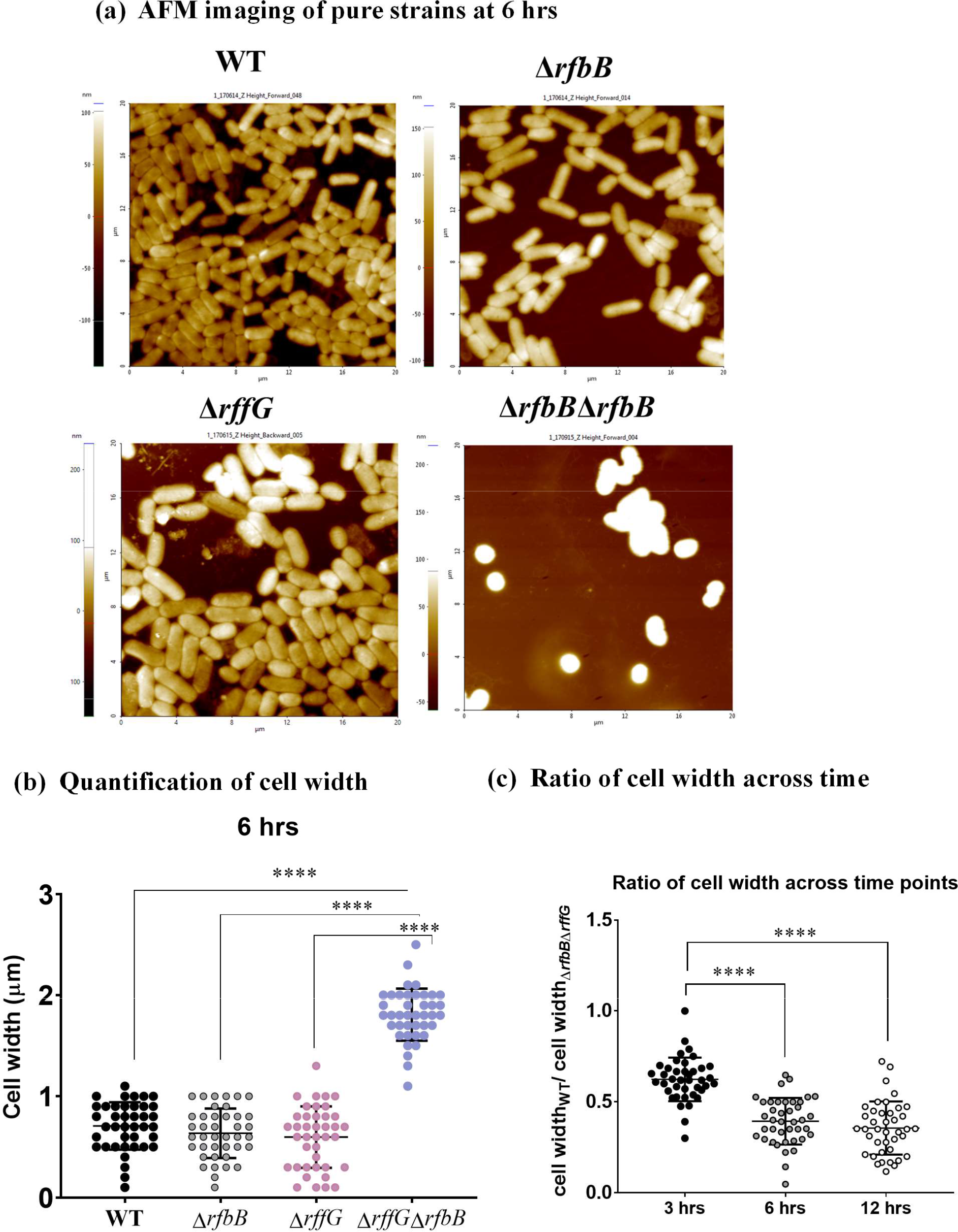
*S*. Typhimurium Δ*rffG*Δ*rfbB* are more rounded in morphology than the WT and single gene-deleted strains. **(a)** WT, Δ*rffG,* Δ*rfbB and* Δ*rffG*Δ*rfbB* strains were grown for 6 hours and AFM images were acquired. **(b)** The cell width of at least 50 cells in each condition were measured and represented after 6 hours of growth in LB broth. **(c)** Ratio of cell width of WT:Δ*rffG*Δ*rfbB* strains estimated across time points. Data are representative of 3 independent experiments and is plotted as mean ± SEM. Statistical analysis was performed using two-way ANOVA, where **** *p* < 0.0001.

### *S*. Typhimurium Δ*rffG*Δ*rfbB* is highly susceptible to bile stress and cell wall targeting antibiotics

Next, we evaluated the susceptibility of these strains to common stresses encountered by *S*. Typhimurium (31). *S.* Typhimurium must withstand a wide range of harsh conditions as it encounters different environments outside the host, within animal intestinal tracts and the intracellular environment of the host phagocytes. In this study, we subjected all the four strains to high temperature (42°C) and monitored the growth for 12 hours. We did not observe any apparent growth defect among the strains (Figure 3a). Furthermore, no differences between strains were observed upon nutrient stress (Figure 3b), different pH (Figure 3c) and osmolar stress (Figure 3d).

**Figure 3.**
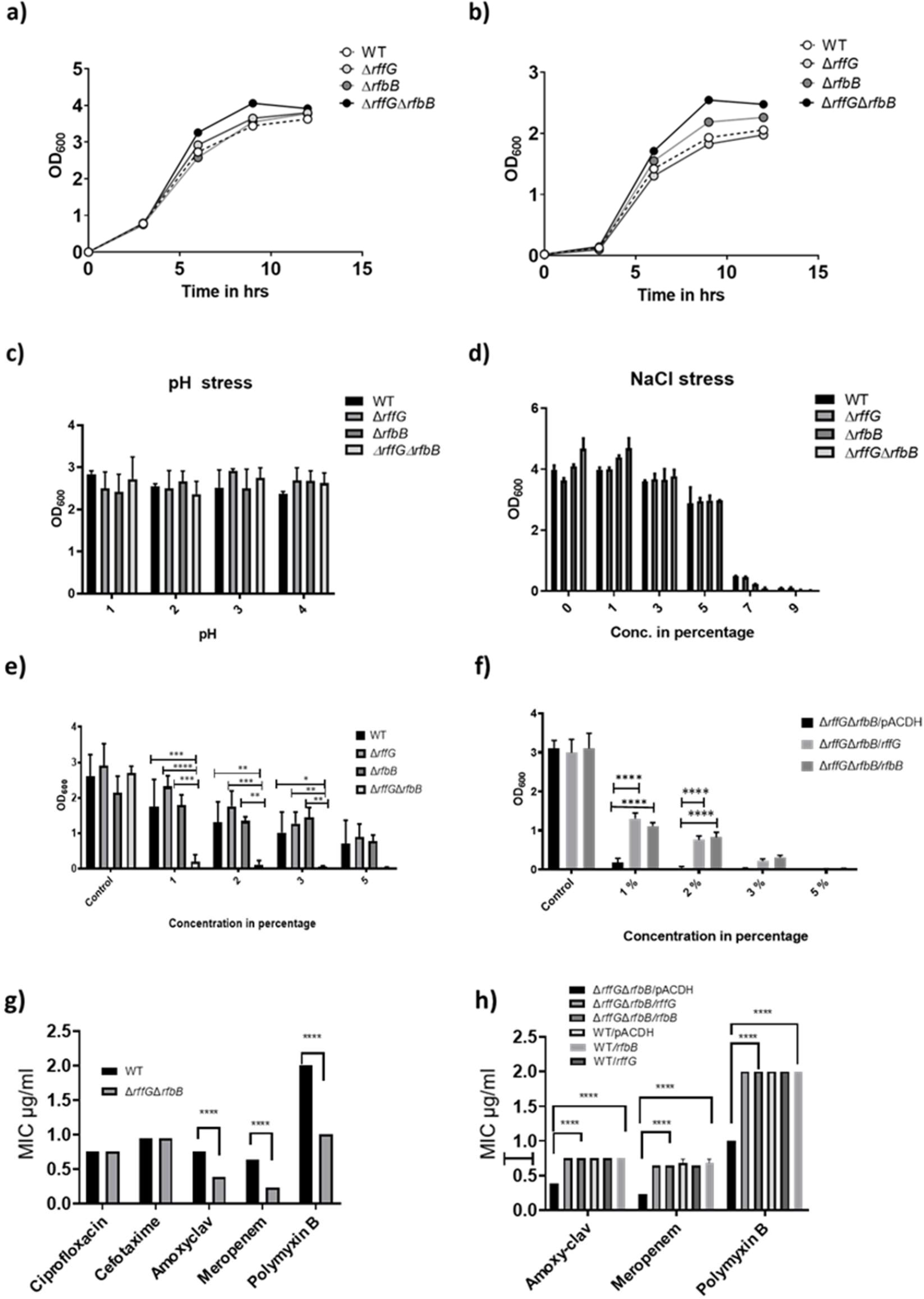
*S*. Typhimurium Δ*rffG*Δ*rfbB* strain is susceptible to bile and cell wall targeting antibiotics. **(a)** *S*. Typhimurium WT, Δ*rffG*, Δ*rfbB* and Δ*rffGΔrfbB* were grown in LB at 42°C and in **(b)** minimal media. Briefly, overnight grown cultures were normalized to 2 O.D. at 600 nm and 0.5% of the respective bacterial cultures were inoculated in an Erlenmeyer flask containing 50 ml LB broth. The flasks were incubated at 37°C under shaking conditions (160 rpm) for 12 hours. At indicated time intervals, 1 ml aliquot of the bacterial cultures were aspirated and the O.D. measured, at 600 nm. **(c)** *S*. Typhimurium WT, Δ*rffG*, Δ*rfbB* and Δ*rffGΔrfbB* growth in 5 ml LB broth of different pH, and **(d)** with different concentrations of NaCl, as indicated. **(e)** *S*. Typhimurium WT, *ΔrffG*, *ΔrfbB* and *ΔrffGΔrfbB* and **(f)** complemented strains, *ΔrffGΔrfbB*/pACDH, *ΔrffGΔrfbB/rffG*, *ΔrffGΔrfbB/rfbB* strains were treated with the indicated concentrations of bile and grown for 8 hours at 37°C and 160 rpm. Bacterial growth was quantified by measuring O.D. at 600 nm. Epsilometer Test (E-Test) results to determine MIC of the **(g)** WT and the Δ*rffG*Δ*rfbB* strains as well as the **(h)** complemented strains. Briefly, overnight grown cultures were normalized to O.D. 2 at 600 nm and plated onto the Mueller Hinton agar plates by spread plate method. Pre-coated Ezy-MIC strips with selected antibiotics were placed in the middle of the agar plates with the help of a sterile swab and the plates were incubated at 37°C for 18 hours. Data are representative of 3 independent experiments and plotted as mean ± SEM. Statistical analysis was performed using two-way ANOVA, where * *p* < 0.05; ** *p* < 0.01; *** *p* < 0.001 and **** *p* < 0.0001.

During the process of colonization of the host, *Salmonella* encounters high concentration of bile in the gall bladder. Pathogens such as *S*. Typhimurium and *S*. Typhi are resistant to high concentration of bile salts (32). Therefore, we examined the susceptibility of these strains by exposing them to different concentrations of bile in the growth medium. However, we did not observe any growth difference between the WT and the single gene-deleted mutants (Figure 3e). Strikingly the Δ*rffG*Δ*rfbB* strain exhibited a significant growth defect under these conditions, and the growth reduction was in a dose-dependent manner. The Δ*rffG*Δ*rfbB* strain was unable to grow at the concentrations of bile greater than 1%. As seen in Figure 3f, the bile-sensitive phenotype of the Δ*rffG*Δ*rfbB* strain could be rescued by complementation with the WT copy of either *rffG* or *rfbB*.

*Salmonella* also encounters a variety of antimicrobial responses primarily driven by the action of different antimicrobial peptides. We studied the antimicrobial susceptibility of these strains by exposing them to commonly used antibiotics. Ciprofloxacin, a fluroquinolone; amoxyclav and meropenem, that are β-lactam antibiotics and polymyxin B, a cationic antimicrobial polypeptide were tested. While there was no notable difference in the minimum inhibitory concentration (MIC) among the strains treated with ciprofloxacin, we observed that the Δ*rffG*Δ*rfbB* strain had lower MIC values for ampicillin, meropenem and polymyxin B (Figure 3g). This phenotype could be restored by complementing the Δ*rffG*Δ*rfbB* strain with the WT copy of either *rffG* or *rfbB in trans* (Figure 3h).

### *S*. Typhimurium Δ*rffG*Δ*rfbB* has a truncated O-antigen profile, defective outer membrane permeability and display auto-aggregation behavior

The susceptibility to cell wall targeting antibiotics and bile indicated that the outer membrane integrity may be compromised in the Δ*rffG*Δ*rfbB* strain. The outer membrane of Gram-negative bacteria contains LPS molecules and OMPs, which constitute the major components of the outer membrane. To investigate any alteration in the outer membrane profile, the LPS and the outer membrane protein (OMP) profile of these strains were analyzed. WT and the single gene-deleted strains did not display any difference in the LPS profile. However, Δ*rffG*Δ*rfbB* strain showed the absence of the repeating O-antigen structure (Figure 4a). The OMP profile of the Δ*rffG*Δ*rfbB* was also monitored and compared to the isogenic WT strain. However, we did not detect any observable difference between the WT and the Δ*rffG*Δ*rfbB* strain (Figure S5). The proteins clustered around 35-45 kDa are likely to be OmpA, OmpD and OmpC (33).

**Figure 4.**
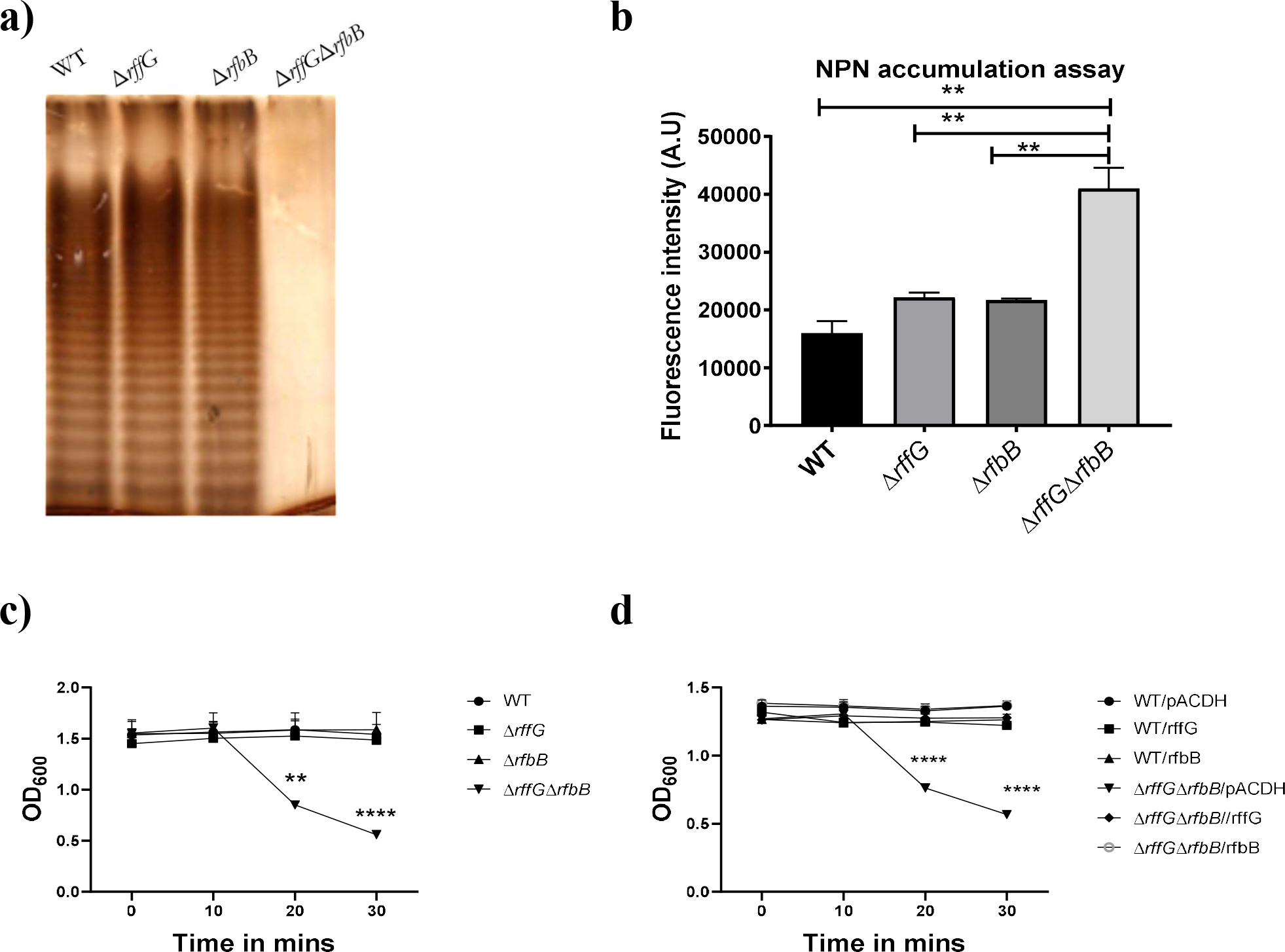
*S*. Typhimurium Δ*rffG*Δ*rfbB* strain displays a truncated O-antigen profile, defective outer membrane permeability and auto-aggregation behavior. **(a)** *S*. Typhimurium WT, Δ*rffG,* Δ*rfbB* and Δ*rffG*Δ*rfbB* strains were grown in LB broth at 37°C for 15 hours. LPS was isolated and resolved on a 12% SDS PAGE. **(b)** *S*. Typhimurium Δ*rffG*Δ*rfbB* accumulates higher amounts of the NPN dye. Briefly, the representative bacterial strains were grown to an O.D. of 0.5 at 600 nm. Cells were harvested, and the bacterial pellet was washed with 5 mM HEPES (pH 7.2) and adjusted to an O.D. of 0.5 at 600 nm. NPN was added to each well at a concentration of 10 μM. Fluorescence excitation and emission were measured at 350 nm and 420 nm, respectively. Statistical analysis for this assay was performed using one-way ANOVA, where ** *p* < 0.01. **(c)** Auto-aggregation behaviour of the WT and the gene-deleted strains. Briefly, 1 ml of 1.5 O.D. normalized, *S*. Typhimurium WT, Δ*rffG*, Δ*rfbB* and Δ*rffG*Δ*rfbB* cultures in LB broth were kept in an upright position for 30 minutes. At every 10-minute interval, 100 μl was aspirated from the top of the solution, and the O.D. was measured at 600 nm. **(d)** Complementation with the WT copy of the gene(s) show a rescue in the phenotype. Data are representative of 3 independent experiments and plotted as mean ± SEM. Statistical analysis was performed using two-way ANOVA, where ** *p* < 0.01 and **** *p* < 0.0001.

The outer membrane of Gram-negative bacteria is unique among all biological membranes by the virtue of its properties to exclude hydrophobic substances. This is due to the presence of LPS on the outer leaflet of the membrane. Thus, the presence of the outer membrane forms a permeability barrier to exclude external agents like hydrophobic antibiotics, detergents, and bile acids (34, 35). The probe 1-N-phenylnaphthylamine (NPN) is used to study biological membranes. It is an environment-sensitive dye, which gives strong fluorescence in phospholipid environments, but fluoresces weakly in aqueous environments (36). Results revealed that the dye accumulated to a similar extent in the WT and the single gene-deleted strains. However, we observed an increased amount of fluorescence in the Δ*rffG*Δ*rfbB* strain (Figure 4b), indicating enhanced basal outer membrane permeability.

During these experiments, and to our surprise, we also observed that the Δ*rffG*Δ*rfbB* strain, if kept undisturbed at room temperature and in a liquid medium form aggregates within a brief period (Figure 4c). This auto-aggregation behavior was optimal at the NaCl concentration of 1%, whereas a modulation in the concentration of NaCl (either higher or lower than 1%) reduced the auto-aggregation rate. We suspect that the auto-aggregation behavior of the Δ*rffG*Δ*rfbB* strain could be due to the absence of O-antigen repeating units which may lead to higher hydrophobicity, culminating in cell-cell aggregation in this strain. The auto-aggregation phenotype of the Δ*rffG*Δ*rfbB* strain was also rescued upon complementation with either *rffG* or *rfbB in trans* (Figure 4d).

### RNA-seq reveals differential gene expression between the WT and the Δ*rffG*Δ*rfbB* strain

Bacterial gene regulation is mediated by a plethora of factors such as transcription factors, nucleoid associated proteins and regulatory small non-coding RNAs. High throughput RNA-sequencing (RNA-Seq) involves sequencing of the complete set of RNA transcripts produced by a cell under any given condition (37). A global transcriptomic analysis of 22 distinct infection-related condition in *S*. Typhimurium was published and presented as a *Salmonella* compendium (38). We investigated the expression of *rffG*, *rfbB* as well as other well-known genes from the *Salmonella* compendium v2.0 (Table S1). Phenotypic characterization of the *S*. Typhimurium WT, Δ*rffG*, Δ*rfbB* and Δ*rffG*Δ*rfbB* strains revealed that major phenotypic differences occurred only in the Δ*rffG*Δ*rfbB* strain, whereas the single gene-deleted strains did not display any observable differences when compared with the isogenic WT strain. To obtain mechanistic insights into the observable phenotypic characteristics of the Δ*rffG*Δ*rfbB,* an RNA-seq analysis was performed in the WT and the Δ*rffG*Δ*rfbB* strains. We chose the early exponential phase to capture the differences in the gene expression pattern, which precede the phenotypic differences that were observed at later time points. RNA-seq analysis revealed considerable gene expression differences between the WT and the Δ*rffG*Δ*rfbB* strain. Those genes which were found to be significantly modulated were mapped to pathways with KEGG Mapper (Figure S6). The results obtained from the RNA-seq analysis were validated by performing functional assays.

Firstly, we observed that a greater number of pathways were downregulated in the Δ*rffG*Δ*rfbB* strain as compared to pathways that were upregulated (Figure S6). We also noted that genes belonging to flagellar assembly pathway as well as genes involved in the infection process, specifically invasion, were among the most significantly downregulated genes in the Δ*rffG*Δ*rfbB* strain. Furthermore, pathways which play an indispensable role during the pathogenesis of *S*. Typhimurium such as chemotaxis as well as quorum sensing were also found to be significantly downmodulated in the Δ*rffG*Δ*rfbB* strain as compared to the isogenic WT strain. In addition, genes involved in the O-antigen biosynthesis and LPS biosynthesis were also found to be downregulated in the Δ*rffG*Δ*rfbB* strain. Altogether, the RNA-seq data highlights that in the Δ*rffG*Δ*rfbB* strain, the pathways which are intricately linked to the virulence of *S*. Typhimurium were greatly suppressed if compared to the WT strain.

Among the pathways which were upregulated in the Δ*rffG*Δ*rfbB* strain as compared to the WT: glycerophospholipid biosynthesis, cationic antimicrobial resistance and nitrogen metabolism were the most prominent and garnered our attention. We reasoned that in the Δ*rffG*Δ*rfbB* strain, the truncated O-antigen structure would result in the remodeling of the outer membrane. It is therefore imperative that genes belonging to the glycerophospholipid metabolism which are known to be involved in the restructuring of the outer membrane would be differentially modulated in the Δ*rffG*Δ*rfbB* strain (39). Since, loss of outer membrane integrity results in more susceptibility towards cationic antimicrobial peptides, the higher expression of the genes which are involved in mediating resistance against the cationic antimicrobial peptide reconfirms and correlates with our earlier observation (40). The upregulation of the genes involved in nitrogen metabolism warrants further investigation. We proceeded to validate the findings obtained from the RNA-seq by performing qRT-PCRs. We focused on the flagellar assembly pathway as well as *Salmonella* infection-related genes which were found to be highly downregulated in the Δ*rffG*Δ*rfbB* strain.

### The flagellar assembly pathway is downregulated and the Δ*rffG*Δ*rfbB* strain displays severe motility defects

Bacterial motility is an important virulence trait of the pathogen *S.* Typhimurium. Findings from the RNA-seq showed that the flagellar assembly pathway was down-regulated in the Δ*rffG*Δ*rfbB* strain (Figure S6). To validate the transcriptomic data, qRT-PCR was performed with a few candidate genes from the flagellar assembly pathway. The flagellar assembly pathway in *S.* Typhimurium is hierarchically organized into three classes. Class I promoter includes *flhDC* which is the master regulator, and its expression is controlled by a plethora of environmental signals. Class II and class III gene products comprise the basal body, hook, filament, and the motor force generator respectively (41, 42). We found that *flhD*, *fliC* and *fljB* were expressed in lower amounts in the single gene-deleted mutants but highly downregulated in the Δ*rffG*Δ*rfbB* strain as compared to the WT strain (Figure 5a-c).

**Figure 5.**
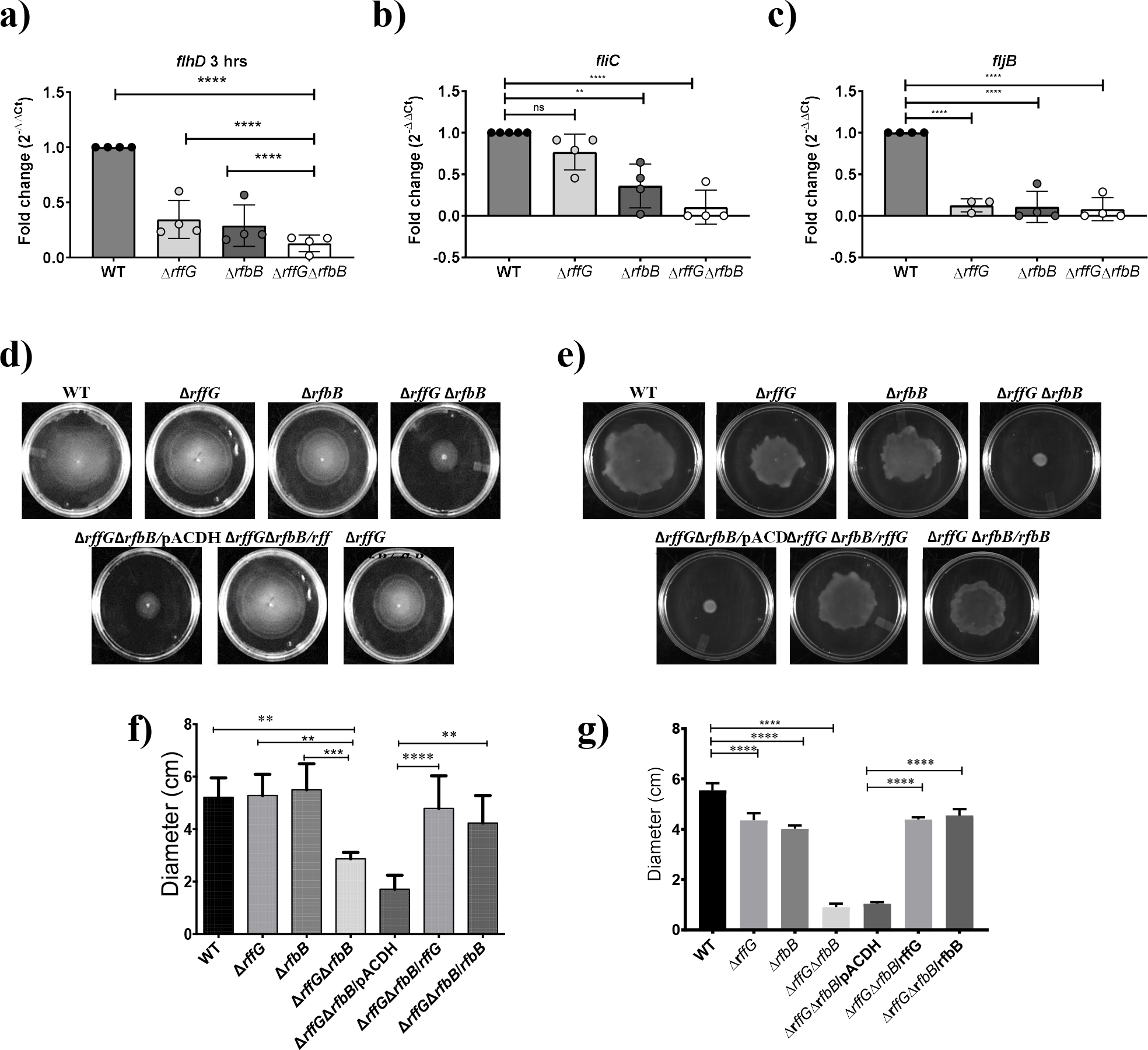
*S*. Typhimurium Δ*rffG*Δ*rfbB* strain is compromised in swimming and swarming motility. Total RNA was extracted from the bacterial samples after 3 hours of growth in LB broth at 37°C and 160 rpm. A qRT-PCR analysis was performed to monitor the expression of the genes, **(a)** *flhD*, **(b)** *fliC* and **(c)** *fljB* and the expression normalized with the reference gene, *gmk*. The **(d) s**wimming and **(e)** swarming motility assays were performed on 0.3% and 0.5% agar, respectively. Equal amounts of cultures were inoculated at the center of the motility-agar plates and the plates were incubated at 37°C for 8 hours in an upright condition. The distance covered by each strain after 8 hours post inoculation for **(f)** swimming and **(g)** swarming motility was measured and analyzed using ImageJ software. Multiple measurements were obtained for each strain. Data are representative of 3 independent experiments, and plotted as mean ± SEM. Statistical analysis was performed using one-way ANOVA, where * *p* < 0.05; ** *p* < 0.01; *** *p* < 0.001 and **** *p* < 0.0001.

To further address whether the transcriptomics data correlates with the actual observation, we performed motility assays to determine the extent of motility defect in these strains. Swimming motility is the result of movement of single cells in aqueous environment. There were no observable differences in swimming motility between the wild type and the single gene-deleted strains (Figure 5d). On the other hand, the Δ*rffG*Δ*rfbB* strain was highly compromised in swimming motility as compared to the WT and the single gene-deleted strains. Complementation with the WT copy of either *rffG* or *rfbB* in Δ*rffG*Δ*rfbB* strain could restore this defect (Figure 5d and Figure 5f). In swarming motility, we observed the single gene-deleted strains to be slightly but significantly compromised as compared to the WT strain. The Δ*rffG*Δ*rfbB* strain, on the other hand, was completely non-motile on swarming agar plates (Figure 5e). Complementation with the WT copy of either *rffG* or *rfbB* in Δ*rffG*Δ*rfbB* strain restored the swarming defect and brought it back to the WT level (Figure 5e and Figure 5g).

### The Δ*rffG*Δ*rfbB* strain displays compromised ability to infect epithelial cells

*S.* Typhimurium employs the products of several genes which are encoded in its genome during infection. Among them, two pathogenicity island related genes are most widely studied. Being an intracellular pathogen, *S.* Typhimurium can infect and persist in the intracellular environment, and this is one of the indispensable features of this pathogen. *S*. Typhimurium can infect and replicate within epithelial cells and macrophages (43). There are three steps in the infection process: adhesion, invasion and intra-cellular replication. Adhesion is a crucial aspect of the infection as it is the first step in establishing contact with the host cell which results in internalization of the pathogen. The invasion of epithelial cells is crucial in maintaining the pathogenicity, persistence, and dissemination to other tissues. *Salmonella* contains *Salmonella* Pathogenicity Island 1 (SPI-1) and *Salmonella* Pathogenicity Island 2 (SPI-2) for invasion and intracellular replication respectively (44, 45).

We had previously determined from the transcriptomic analysis that genes belonging to the SPI-1 pathway were downregulated in the Δ*rffG*Δ*rfbB* strain. To validate these findings, qRT-PCRs of some of the key genes from the SPI-1 pathway were performed. We found that the expression of *hilA, hilD* and *sipC*, all of which regulate the expression of other key downstream effectors of the SPI-1 pathway, was two-fold downregulated in the Δ*rffG*Δ*rfbB* as compared to the WT strain (Figure 6a-c). We also noted that *hilA*, *hilD* and *sipC* was downregulated in the single gene-deleted strains as compared to the WT strain. Overall, our qRT-PCR data agrees with the transcriptomics data.

**Figure 6.**
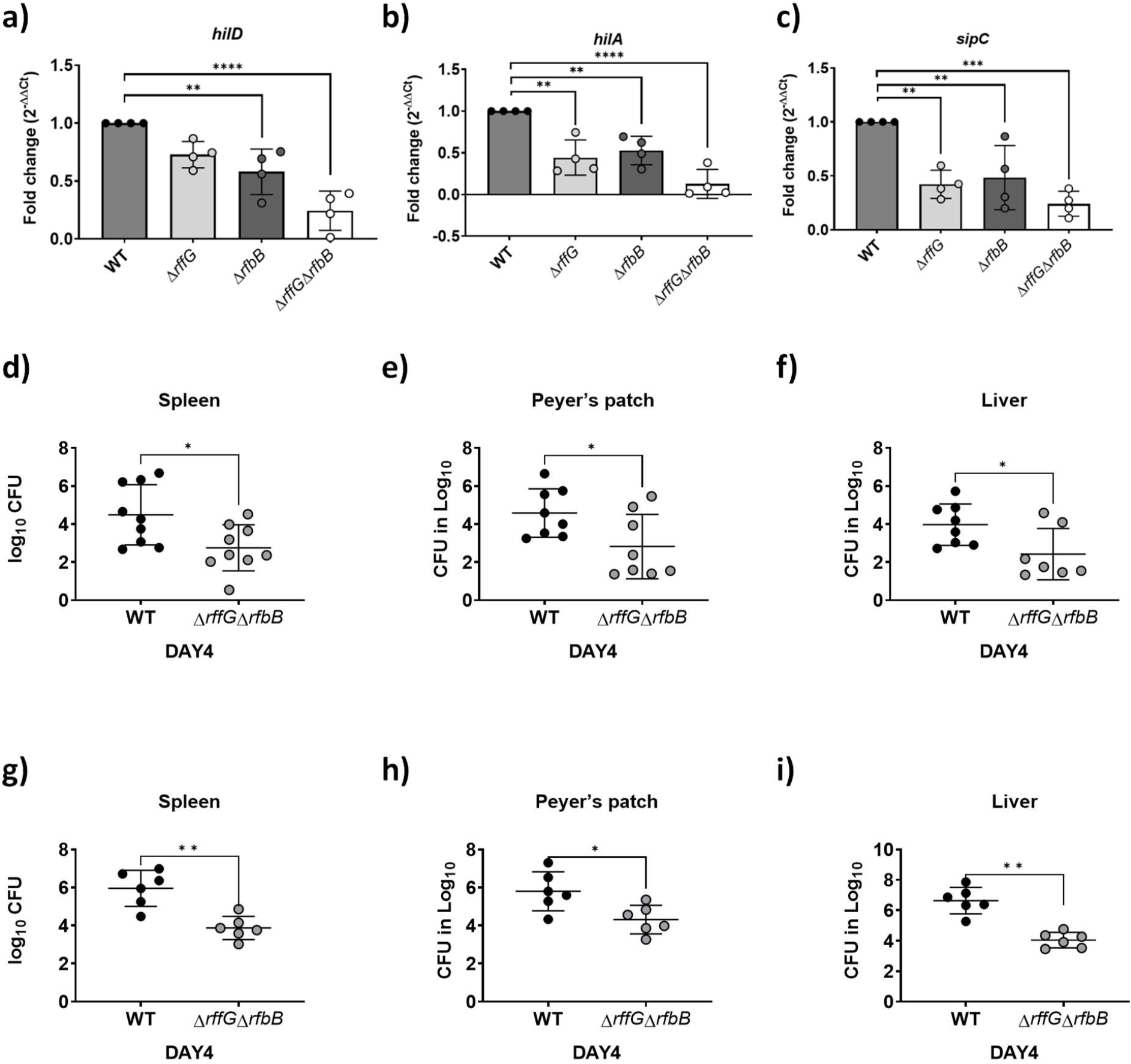
*S.* Typhimurium Δ*rffG*Δ*rfbB* strain is less proficient in colonizing different organs. Total RNA was extracted from the bacterial samples after 3 hours of growth in LB broth at 37°C and 160 rpm. A qRT-PCR analysis was performed to monitor the expression of the SPI-1 genes, **(a)** *hilD* **(b)** *hilA* and **(c)** *sipC,* and the expression normalized with the reference gene, *gmk.* Bacterial burden in the organs was estimated after **(d-f)** oral and **(g-i)** intraperitoneal infection of C57BL/6 mice with the WT and the Δ*rffG*Δ*rfbB* strains. Infected mice were sacrificed on day 4 post-infection, organs were harvested, and the bacterial burden in the spleen, Peyer’s Patch and the liver was estimated by plating serial dilutions of the tissue homogenate on LB agar plate. Gene expression analysis data are representative of 3 independent experiments plotted as mean ± SEM and statistical analysis was performed using one-way ANOVA, where * *p* < 0.05; ** *p* < 0.01; *** *p* < 0.001 and **** *p* < 0.0001. For estimating bacterial burden in organs, statistical analysis was performed using unpaired t-test.

One of the critical steps in the infection process of *S*. Typhimurium is the establishment of successful infection niche in the intestinal epithelial cells. Breaching the epithelial barrier is paramount in dissemination of the pathogen to different organs. We studied the epithelial infection process by using HeLa cells as a model system. We studied all the three steps in the active infection processes in HeLa cells (Figure S7a-c). We observed that the single gene-deleted strains were slightly more compromised than the WT in the adhesion to the HeLa cells, whereas the Δ*rffG*Δ*rfbB* strain was significantly more compromised than the other strains with respect to adhesion. Additionally, the number of bacteria that successfully invaded the HeLa cells were measured by lysing the cells and counting the number of bacteria obtained at 2 hours post infection. We found that the single gene-deleted mutants, Δ*rffG* and Δ*rfbB* also showed a slight, although significant reduction in the number of viable bacteria at 2 hours post infection when compared to the WT strain. Interestingly, the Δ*rffG*Δ*rfbB* strain showed a significant reduction of the number of viable bacteria at 2 hours post infection. Furthermore, we studied the ability of the bacteria which have invaded the HeLa cells to replicate in the intracellular environment. To this end, we obtained and enumerated the number of viable bacteria at 18 hours post infection. We did not observe any significant difference in the fold change in CFU between 2 hours and 18 hours among the different strains. Altogether, our results indicate that the Δ*rffG*Δ*rfbB* strain is compromised in comparison to the other strains in its ability to adhere and subsequently invade the epithelial cells but does not show any defect in the intracellular replication.

### The Δ*rffG*Δ*rfbB* strain is less proficient in colonizing different organs and induces lower pro-inflammatory cytokine response

We had observed that the flagellar pathway as well as the SPI-1 pathway is down modulated in the Δ*rffG*Δ*rfbB* strain. We were interested in determining whether it would have any effect on the organ colonization in mice. Hence, we proceeded to infect C57BL/6 mice through oral as well as intraperitoneal routes and determine the organ CFU burden at 4-day post infection (Figure 6d-i). In the oral infection model, the pathogen must traverse the epithelial barrier, whereas in the intra-peritoneal (i.p.) infection model, the pathogen directly enters the systemic circulation. We infected C57BL/6 mice orally with 1×10^8^ CFU of the WT and the Δ*rffG*Δ*rfbB* strain. At four-day post infection, we found that the Δ*rffG*Δ*rfbB* strain displayed two log-fold lower infection burden in the Peyer’s patches, liver, and spleen (Figure 6d-f). It is possible that the lowered CFU burden observed in the different organs might be due to the reduced expression of SPI-1 genes needed to traverse the epithelial barrier. To address this question, C57BL/6 mice were intraperitoneally infected with the WT and the Δ*rffG*Δ*rfbB* strains as intraperitoneal infection would not require active breach of the epithelial barrier. At four-day post infection through the intraperitoneal route, we found that the CFU burden of the Δ*rffG*Δ*rfbB* strain was two log-folds lower as compared to the WT (Figure 6g-h). This clearly suggested that the low proficiency to infect and colonize different organs was not solely due to the reduced expression of the SPI-1 genes in the Δ*rffG*Δ*rfbB* strain but could be due to multiple and varied mechanisms at play. Subsequently, we investigated whether the Δ*rffG*Δ*rfbB* strain had the ability to induce a pro-inflammatory cytokine response in the host. To address this question, we collected serum from the WT and Δ*rffG*Δ*rfbB* infected mice at four-day post infection. We found that the Δ*rffG*Δ*rfbB* strain induced lesser pro-inflammatory cytokines such as TNF-α, IL-6 and IFN-γ than the WT strain in both the oral (Figure 7a-c) and the i.p. (Figure 7d-f) modes of infection.

**Figure 7.**
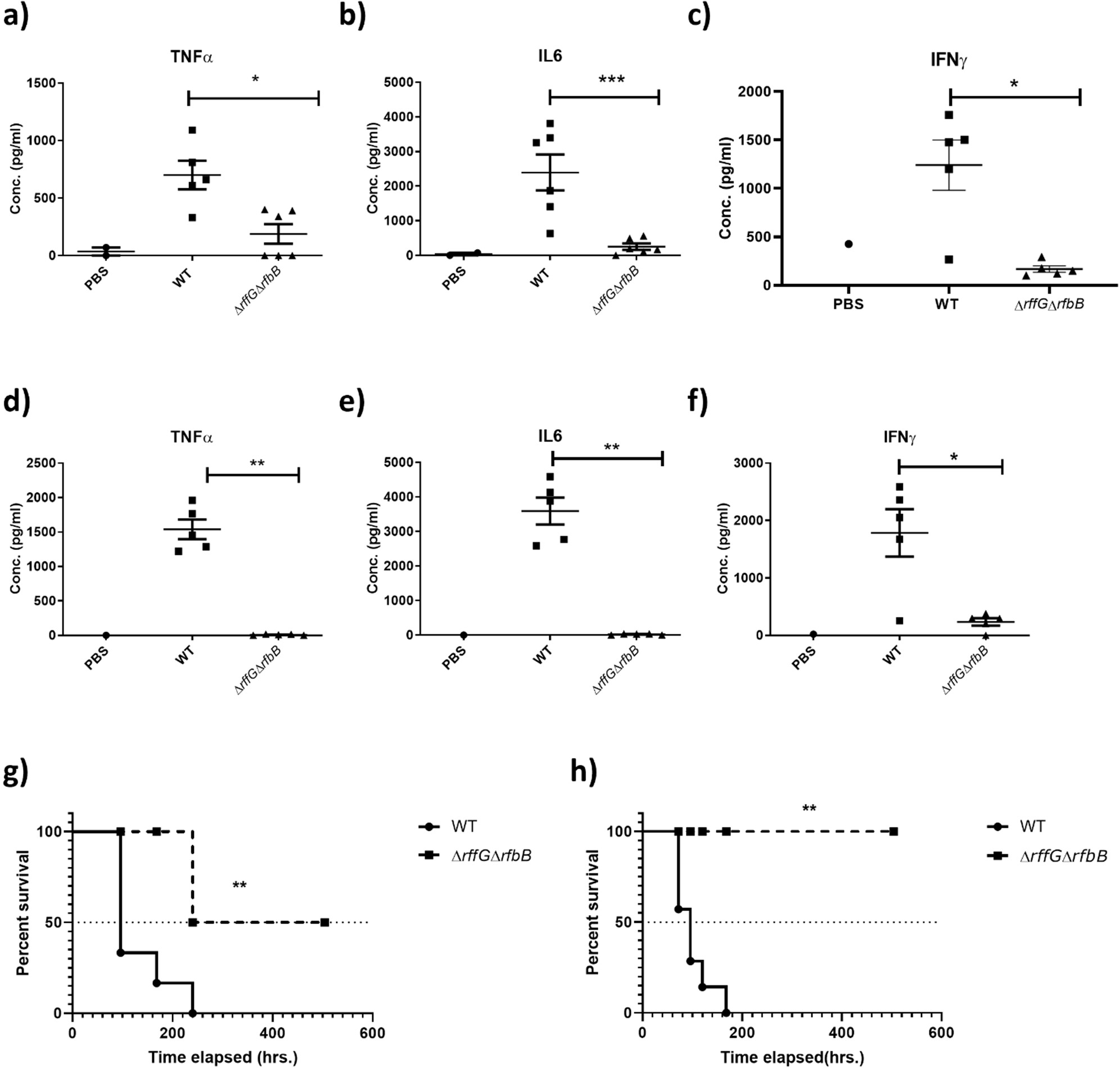
*S*. Typhimurium Δ*rff*GΔ*rfb*B strain is highly attenuated in oral and intraperitoneal mice models of infection. The levels of pro-inflammatory cytokines **(a)** TNFα **(b)** IL6, and **(c)** IFNγ at day 4 post-infection were estimated from the serum collected from mice infected orally with 1×10^8^ CFU of the WT and Δ*rff*GΔ*rfb*B strains. Similarly, mice infected intraperitoneally with 1×10^3^ CFU of *S*. Typhimurium WT and Δ*rff*GΔ*rfb*B were sacrificed at day 4 post-infection and the serum levels of cytokines, **(d)** TNFα **(e)** IL6 and **(f)** IFNγ were quantified. Kaplan-Meier survival analysis comparing mice survival for 21 days upon infection with the WT and the Δ*rffG*Δ*rfbB* strains, infected either, **(g)** orally or **(h)** intraperitoneally. Data are representative of 3 independent experiments plotted as mean ± SEM. Statistical analysis was performed using Log-rank (Mantel-Cox) test, where ** *p* < 0.005 and *** *p* < 0.0001. For cytokine estimation, statistical analysis was performed using unpaired t-test, where * *p* < 0.05; ** *p* < 0.01 and *** *p* < 0.001.

### *ΔrffGΔrfbB* strain is attenuated in both oral and intraperitoneal model of *S*. Typhimurium infection

Finally, we monitored the survival of the mice after oral and intraperitoneal infection for 21 days post infection (Figure 7). Mice infected with the WT strain succumbed to infection as early as 4 days post infection and 100% mortality was observed by day 12. In contrast, infection with the Δ*rffG*Δ*rfbB* strain resulted in prolonged mice survival. By day 21, only 20% mortality was observed (Figure 7g). Similarly, in the i.p. infected mice all the WT-infected mice succumbed to the infection by day 7 post infection whereas the Δ*rffG*Δ*rfbB* strain showed no mortality till 21 days post infection (Figure 7h). Overall, the Δ*rffG*Δ*rfbB* strain was highly attenuated in causing systemic disease in mice compared to the WT strain. Although the Δ*rffG*Δ*rfbB* strain had the ability to colonize different organs, it was less proficient in the induction of pro-inflammatory cytokine response and did not cause lethal infection when administered via oral as well as the i.p. routes of infection.

## Discussion

In this study, we sought to investigate how the absence of the genes, *rffG* and *rfbB* affect the physiology of *S*. Typhimurium. The WT strain and the single gene-deleted strains displayed no apparent growth differences in LB broth, whereas the Δ*rffG*Δ*rfbB* strain demonstrated a considerable lag phase. The Δ*rffG*Δ*rfbB* strain also showed a higher OD in the stationary phase as compared to the other strains but did not display any difference in the CFU (Figure S3). The OD of a solution is the property of the concentration of the solutes as well as the size and shape of the solute particles (https://www.nature.com/articles/srep38828). The Δ*rffG*Δ*rfbB* strain exhibits greater width as compared to the other strains. It is possible that the change in cell shape in the double gene-deleted strain could account for an increased O.D. when compared to the WT or single gene-deleted strains (46, 47).

The AFM studies revealed that the Δ*rffG*Δ*rfbB* strain displayed a predominantly round morphology as compared to WT and the single gene-deleted strains (Figure 2a and Figure S4). We also noted that the Δ*rffG*Δ*rfbB* strain had greater cellular width when compared with the other strains (Figure 2b). Interestingly, *Salmonella* mutants lacking ECA (48) or LPS (49) do not show the round phenotype as seen in Δ*rffG*Δ*rfbB* strain. Cell morphology is determined by the arrangement of the cytoskeletal proteins and peptidoglycan synthesis. Previous reports suggest that the mutations in *mre* and *rod*, which are components of cytoskeletal arrangements in *S*. Typhimurium gives rise to round cellular morphology (50). Also, peptidoglycan biosynthesis is important in maintaining the cellular shape in *S*. Typhimurium (51). The accumulation of metabolic intermediates of the O-antigen and ECA biosynthesis has also been reported to affect the cell shape (52). MinC and MinD are required for proper cell division. In the absence of MinC or MinD, the FtsZ ring fails to locate to the middle of a cell, leading to abnormal cell division and morphological aberrations. Multiple FtsZ rings are randomly and simultaneously formed in various positions of a cell, which results in the bacterial cell separating into more than two daughter cells (53). The effect of downregulation of these two genes in Δ*rffG*Δ*rfbB* strain needs to be further investigated. Further work is needed to shed light on the mechanisms at play which affects the cell morphology in the Δ*rffG*Δ*rfbB* strain. Another interesting observation was the size of the colonies on agar plates formed by the Δ*rffG*Δ*rfbB* strain was smaller in comparison to the WT and the single gene-deleted mutants (Figure 1a-c). Earlier studies have reported the conditional lethality of the round cell mutants on the solid versus the liquid medium. It was reported that the round cell mutants of *S*. Typhimurium were more susceptible to growth on solid medium than in liquid medium (54). This observation was correlated to an anomalous cell division process in *rodA* and *mre* mutants on solid medium and not due to their envelop characteristics. Thus, it is possible that while growing on the solid medium, like LB agar, the Δ*rffG*Δ*rfbB* strain encounters certain stresses which leads to a reduction in growth as opposed to growth in the liquid medium. These studies highlight the fact that the growth requirements for a strain may not be the same in solid and liquid media.

We had also subjected these strains to various stresses that *S*. Typhimurium commonly encounters during the host colonization process. We detected no growth difference in the osmolarity-induced stress and pH-induced stress among the strains (Figure 3c-d). However, the Δ*rffG*Δ*rfbB* strain was found to be highly susceptible to bile-induced stress (Figure 3e-f). LPS acts as a membrane barrier to bile salt uptake (55) and this observation is in accordance with previous studies that showed that LPS mutants are susceptible to bile (56). *Salmonella* strains lacking ECA do not show any altered morphology, LPS profile or motility, although they are attenuated in mice models of infection (57). In fact, it is possible that the reduced virulence is because *Salmonella* strains lacking ECA are sensitive to bile (48). We also found that the Δ*rffG*Δ*rfbB* strain showed lower MIC values against cell wall targeting antibiotics such as amoxiclav, meropenem and polymyxin B (Figure 3g). Upon complementation, the MIC values of the Δ*rffG*Δ*rfbB* strain was approximately equal to the WT strain (Figure 3h). Since the Δ*rffG*Δ*rfbB* strain has a truncated LPS (Figure 4a) which lacks the O-antigen and the ECA, these may change the properties of the outer membrane which could, in turn lead to greater susceptibility to the cell wall targeting antibiotics and cationic polypeptides. To address this, we performed NPN dye accumulation assay to check for the outer membrane permeability of the four strains (Figure 4b). Our data demonstrated that the outer membrane permeability was indeed significantly increased in the Δ*rffG*Δ*rfbB* strain.

Moreover, we had observed during our investigations that the Δ*rffG*Δ*rfbB* strain has a property to form auto aggregates in a static liquid medium (Figure 4c). Auto-aggregation was found to be dependent on the concentration of salts in the medium (data not shown) and the phenotype could be rescued upon gene complementation (Figure 4d). Earlier reports have implicated the overexpression of autotransporter adhesins and FimH during auto aggregation in liquid cultures of *E. coli* (58, 59). Overexpression of *misL*, a fimbriae adhesin from *S.* Typhimurium, in *E. coli* AAEC189Δ*flu*, a strain that lacks fimbriae, leads to bacterial aggregation and settling (60). Furthermore, the composition of the outer membrane proteins also determines the ability of the cells to auto aggregate. However, as we did not detect any major differences in the OMP profile in our strains (Figure S5), we could rule out the fact that the property of auto aggregation was mediated by the upregulation of the autotransporters. Most likely, the explanation for enhanced auto aggregation may be due to the increased membrane hydrophobicity in the Δ*rffG*Δ*rfbB* strain (Figure 4b) due to the absence of the hydrophilic O-antigen repeating structures. Together these findings revealed that the Δ*rffG*Δ*rfbB* strain showed distinct phenotypic characteristics when compared to the isogenic WT and the single gene-deleted strains.

Subsequently, we studied the differences at transcriptional level to understand how these phenotypic differences could influence other physiological functions which are fundamental to pathogenesis of *S*. Typhimurium. Based on our prior observations from the phenotypes in the wild type and the single gene-deleted strains, Δ*rffG* and Δ*rfbB*, we determined that the gene products are mutually exchangeable, and one can complement each other’s function. This led us to speculate that, at the transcript level, the major differences would occur between the WT and the Δ*rffG*Δ*rfbB* strains. Hence, we performed a transcriptome analysis (RNA-seq) of the WT and the Δ*rffG*Δ*rfbB* strains and observed major alterations in the pathways involved in motility, invasion, chemotaxis, etc. (Figure S6). We proceeded to validate the observed differences in the RNA-seq data with actual physiological processes in *S*. Typhimurium. The flagellar assembly pathway was one among the prominent pathways that was highly downregulated in the Δ*rffG*Δ*rfbB* strain and we performed motility assay with all the four strains. *S.* Typhimurium exhibits both swarming and swimming motility. In swarming motility, the cells move on semisolid surface as cell rafts embedded in a complex ‘slime’ which is composed of polysaccharide, proteins, surfactants etc. In contrast, swimming motility is the result of movement of single cells in aqueous environment (61). The qRT-PCR revealed differences in the expression of flagellar assembly genes among these strains (Figure 5a-c). Motility experiments revealed that the Δ*rffG*Δ*rfbB* strain was highly compromised in the swimming motility as compared to the WT and the single gene-deleted strains (Figure 5d and Figure 5f). However, in swarming motility the Δ*rffG*Δ*rfbB* strain was completely non motile (Figure 5e and Figure 5g). Single gene-deleted strains also showed a slight but significant defect in swarming motility as compared to the isogenic WT strain. Swarmer cells are hyperflagellated and morphologically different than their swimming counterparts. Multiple studies have reported the altered motility pattern in the LPS truncated mutants (62). Several explanations have been put forth, such as the O-antigen of the LPS serves as wettability factor which facilitates the movement of the bacteria on surfaces during swarming motility (63). Also, truncation of the LPS molecule leads to the activation of outer membrane stress response pathway which is involved in the repression of motility though the down regulation of the flagellar assembly pathway. In *E. coli*, the Rcs two component signaling pathway has also been shown to play a role in repression of motility in LPS mutants (64). Lack of proper LPS has been known to perturb the outer membrane leading to activation of signaling pathways, including RpoE and Rcs phosphor relay system. In turn, this leads to the degradation of FlhDC, the class I regulator, leading to downregulation of flagellar synthesis and, consequently, reduced motility (5). It is possible that in the Δ*rffG*Δ*rfbB* strain the absence of the O-antigen and ECA leads to loss of surface lubrication and the activation of the outer membrane stress response pathway, which leads to the downregulation of flagellar biosynthesis resulting in the complete loss of swarming and highly compromised swimming behavior.

Another, pathway that was found to be highly downmodulated in the Δ*rffG*Δ*rfbB* strain as compared to the WT was the SPI-1 pathway (Figure 6a-c). Initially, we found that there was a small but significant difference in the invasion ability between the WT and the single gene-deleted strains in HeLa cells. However, the Δ*rffG*Δ*rfbB* strain was found to be highly compromised in its ability for invasion (Figure S7). The significant difference in the invasion for the Δ*rffG*Δ*rfbB* strain observed in epithelial cells like HeLa, could possibly be because *Salmonella* has to employ the action of SPI-1 effector proteins to invade these cells (65). Some studies have shown that the SPI-1 pathway and the flagellar pathway are co-regulated (66). The repression of the flagellar pathway has been shown to be mediated by the activation of outer membrane stress response system (5). Also, it is interesting to note that both these pathways are downregulated during the intracellular replication (67). It would be interesting to investigate whether the outer membrane stress response pathway experienced by *S*. Typhimurium during intracellular replication is responsible for the downregulation of the flagellar and the SPI-1 pathways.

Both SPI-1 and the flagellar biosynthesis pathway are essential for successful colonization of mice (68, 69). Studies have shown that *inv* mutants of *S*. Typhimurium, which do not have functional SPI-1, when administered perorally to BALB/c mice, resulted in higher 50% lethal doses (LD_50_) compared to their WT parent strains. However, there were no such differences in the observed LD_50_ when these strains were administered intraperitoneally. In addition, *inv* mutants were compromised in their ability to colonize the Peyer’s patches, small intestinal wall, and the spleen when administered perorally, although when administered intraperitoneally, they showed no difference in their ability to colonize the spleen (70). Here, we further analyzed the infection potential of the Δ*rffG*Δ*rfbB* strain. We observed that the Δ*rffG*Δ*rfbB* was more compromised when compared to the WT strain in colonizing the peripheral organs such as liver, spleen and Peyer’s patches (Figure 6d-i). Also, the Δ*rffG*Δ*rfbB* strain was found to induce lower pro-inflammatory cytokine response in mice (Figure 7a-f). This observation is consistent with previous findings in *Candida albicans*, where it was observed that a GAL102p mutant (a homolog of dTDP-glucose 4,6-dehydratase) was unable to grow in resident peritoneal macrophages. The mutant also elicited a weak pro-inflammatory cytokine response *in vitro* as well as in an *in vivo* mouse model of systemic candidiasis. While the pro-inflammatory responses were found to lowered in this infection model, the GAL102p mutant however had demonstrated an enhanced anti-inflammatory response depicted by serum IL4 measurements (19). Both O-antigen, ECA and the flagella are important antigens recognized by the immune system (71, 72). The absence or down regulation of these components would cause aberrant immune responses in mice infected with the Δ*rffG*Δ*rfbB* strain. Consistent with our previous findings, we observed that the mice infected with the Δ*rffG*Δ*rfbB* strain did not succumb to infection even after 21 days (Figure 7g-h). Earlier studies have also reported that the LPS or ECA mutant are highly attenuated in *S.* Typhimurium oral infection model (3, 4). It is possible that in the absence of a fully operational SPI-1 and motility pathway, the infection cycles during infection are non-optimal. Also, the absence of the O-antigen and ECA in the Δ*rffG*Δ*rfbB* strain may make these strains more susceptible to complement mediated lysis (73). Thereby it is likely that the host can eventually clear the infection when infected with the Δ*rffG*Δ*rfbB* strain. Overall, the absence of the O-antigen, ECA, down regulation of SPI-1 and flagellar biosynthesis pathway could result in a highly attenuated phenotype with respect to virulence as observed in the Δ*rffG*Δ*rfbB* strain.

We present a model that accounts for the key observations (Figure 8). Briefly, the genes, *rffG* and *rfbB* are paralogs that encode the enzyme dTDP-glucose 4,6-dehydratase, which is known to catalyze the intermittent steps of O-antigen and ECA biosynthesis. Consequently, functional loss of dTDP-glucose 4,6-dehydratase renders *S*. Typhimurium strains incapable of synthesizing both O-antigen and ECA. This leads to profound physiological differences between the *S*. Typhimurium, WT and the Δ*rffG*Δ*rfbB* strains. In this study, we report that the Δ*rffG*Δ*rfbB* strain exhibits a distinct cellular morphology, altered LPS profile, increased outer membrane permeability and susceptibility to toxic chemicals. Functional loss of dTDP-glucose 4,6-dehydratase also resulted in inhibition of motility and a reduced ability to colonize different organs in the two distinct and established models of infection in mice. Consequently, the Δ*rffG*Δ*rfbB* strain displayed significant virulence attenuation as compared to the WT. The LPS biosynthesis pathway has recently generated a lot of optimism as a drug target candidate (74). Also, the LPS and ECA mutants are constantly being evaluated as vaccine candidates. To the best of our knowledge, this is one of the primary reports to demonstrate in detail the physiological consequences of the loss of both O-antigen and ECA and the associated gene expression pattern differences in an *S*. Typhimurium strain. This study, therefore, lays the foundation for further research that can lead to a better understanding of the functional consequences of the loss of outer membrane components such as O-antigen and ECA antigen and its effect on *S*. Typhimurium physiology and pathogenesis.

**Figure 8.**
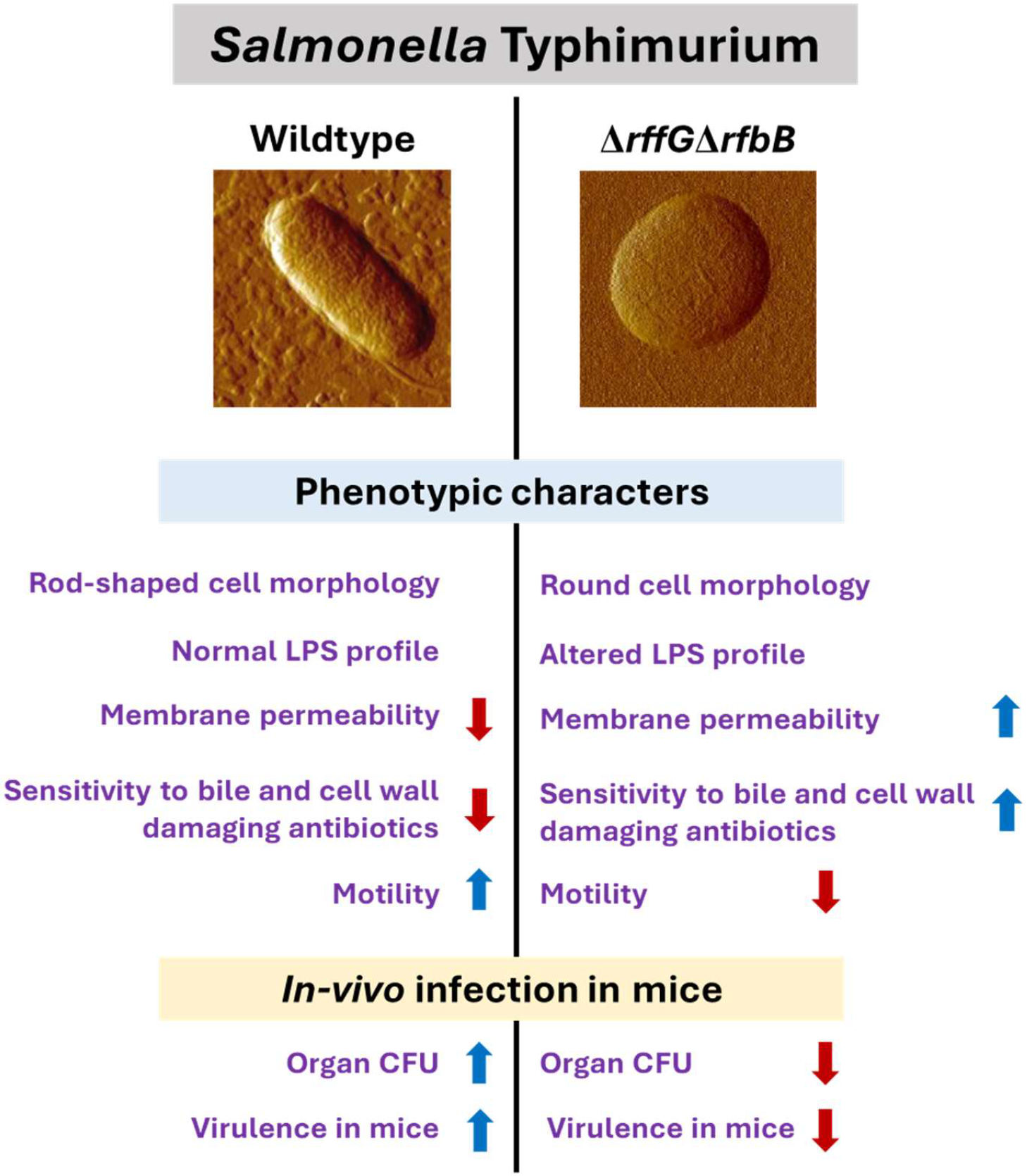
*S*. Typhimurium WT and Δ*rffG*Δ*rfbB* show distinct physiological and phenotypic differences. The genes, *rffG* and *rfbB,* encode the enzyme dTDP-glucose 4,6-dehydratase, which catalyzes intermittent steps of O-antigen and ECA biosynthesis. Functional loss of dTDP-glucose 4,6-dehydratase renders *S.* Typhimurium strains incapable of synthesizing both O-antigen and ECA. This leads to profound physiological differences between the *S.* Typhimurium, WT and the Δ*rffG*Δ*rfbB* strains. The Δ*rffG*Δ*rfbB* strain exhibits distinct cellular morphology, altered LPS profile, increased outer membrane permeability and susceptibility to bile and cell wall targeting antibiotics. Functional loss of dTDP-glucose 4,6-dehydratase also led to inhibition of motility and a reduced ability to colonize different organs in the mouse model of infection. Consequently, the Δ*rffG*Δ*rfbB* strain displayed significant virulence attenuation as compared to the WT.

## Materials and Methods

### Bacterial strains and growth conditions

The bacterial strains used in this study are listed in Table S2. All bacterial cultures were grown in Luria-Bertani (LB) medium consisting of 10 g/L tryptone (HiMedia Laboratories, Mumbai, India), 10 g/L NaCl (Merck, Darmstadt, Germany), and 5 g/L yeast extract (HiMedia Laboratories) at 37°C. Strains containing pKD46 were cultured at 30°C with constant shaking at 160 rpm. Antibiotics were used at the following concentrations: kanamycin, 50μg/ml; chloramphenicol, 30μg/ml; tetracycline, 10μg/ml and/or ampicillin, 100μg/ml.

### Generation of knockout strains and complementation

All single gene-deleted strains used in this study were generated using the one-step gene disruption strategy (29). *S*. Typhimurium 14028s (WT) was used as the parent strain for all experiments, unless otherwise mentioned. Briefly, for the construction of Δ*rffG*, primers listed in Table S3 (Sigma, Bangalore, India) were designed to amplify the kanamycin cassette from the template, pKD4, using PCR. The resulting PCR product was purified and electroporated into the WT strain harboring pKD46, which expresses the λ Red recombinase. A similar methodology was followed for generating the Δ*rfbB* strain, where the gene was replaced with the chloramphenicol cassette (pKD3 template). The double gene-deleted strain, Δ*rffG*Δ*rfbB* was constructed by amplifying the region of the gene from the single knockout strain and electroporating the amplicon into Δ*rffG* strain harboring pKD46. All the mutants were confirmed by PCR amplification using primers designed to anneal ∼100 bp upstream and downstream of the gene (Table S3).

### Cloning of genes for trans-complementation

*S*. Typhimurium 14028s genomic DNA was used as a template for the PCR amplification of genes *rffG* and *rfbB* with the specific primers (Table S3) using the Phusion DNA polymerase. The genes, *rffG* and *rfbB* were cloned between the sites, NcoI and HindIII in the pACDH plasmid. Positive clones were confirmed by Sanger sequencing (Aggrigenome, India). The positive clones and the control vector pACDH were then transformed into *S.* Typhimurium WT and Δ*rffG*Δ*rfbB* by electroporation to generate the complemented strains.

### Stress assays

*S*. Typhimurium WT, Δ*rfbB*, Δ*rffG* and Δ*rfbB*Δ*rffG* were grown in 3 ml LB or LB with appropriate concentration of antibiotics overnight at 37°C with 160 rpm. The OD of the pre-inoculum was normalised to 2.0, at 600 nm and 50μl of the culture was added to 5ml of LB alone or with different concentration of either bile salts (Sigma Aldrich, USA) (1%, 2%, 3% and 5%), NaCl (1%, 3%, 5%, 7% and 9%) or pH range. The cultures were incubated at 37°C under shaking conditions with shaking at 160 rpm for 8 hours and absorbance at 600 nm were measured subsequently in an UV-Visible spectrophotometer (Shimadzu) (33).

### MIC determination

The MIC determination values was performed by E-tests using Ezy-MIC strips (HiMedia, Mumbai, India) according to the manufacturer’s protocol. Briefly, the overnight grown cultures were normalised to 2.0 OD at 600nm and spread-plated onto the Mueller Hinton agar plates by spread plate method. Precoated MIC strips with antibiotics were placed at the middle of the agar plates with the help of a sterile swab and the plates were kept at 37°C for 18 hours. The MIC was read at the point of the strip at which zone of clearance coincided with the strip (75).

### NPN assay

The outer membrane permeability was measured with the help of NPN assay as previously described (5). Briefly, the cells were grown till OD 0.5 at 600 nm, the cells were harvested by centrifugation. The pellet was washed with 5mM HEPES (pH 7.2) and adjusted to an OD of 0.5 at 600 nm. NPN was added to each well at the final concentration of 10μM. 200μl was added to a flat bottom 96-well plate and fluorescence excitation and emission was measured at 350 nm and 420 nm, respectively.

### Analysis of the LPS profile

Bacterial cells were grown for 16 hours samples and the OD was normalised to 2.0 OD at 600 nm. Cells were centrifuged and the pellets were suspended in 150 ml of lysis buffer containing proteinase K (Thermo Scientific, Walter, MA, USA) followed by hot phenol extraction and a subsequent extraction of the aqueous phase with diethyl ether. LPS was separated on 12% (w/v) acrylamide gels using a Tricine-SDS buffer system and visualized by silver staining (76).

### Auto-aggregation assay

The bacterial growth for every strain was normalised to OD 2.0 at 600 nm and the cultures were kept in a static, upright position for 30 minutes. At regular intervals of 10 minutes, 100μl of the culture was collected from the top and the absorbance was measured at 600 nm. Absorbance at 600 nm versus time was plotted over a 30-minute time interval (58).

### RNA isolation and cDNA synthesis

Total RNA was isolated from the cells grown for 3 hours in LB at 37°C. For total RNA preparation, bacterial cultures grown for the mentioned time points (as indicated in the figure), pelleted and stabilized with RNAprotect Bacteria Reagent (Qiagen) according to the protocol recommended by the manufacturer and stored at -80°C, until further use. Total RNA was extracted using the TRIzol reagent (Sigma, St. Louise, Missouri, USA) as per the manufacturer’s instructions. The RNA pellet obtained, was dried and dissolved in 15μl of DEPC water. The purity and concentration of RNA was measured using Nanodrop spectrophotometer (Thermo Scientific, Waltham, MA, USA). Genomic DNA was removed using the RNase-free DNase I (Thermo Fisher Scientific). Subsequently, 2-5μg DNase-treated RNA was reversed transcribed to cDNA using the RevertAid First Strand cDNA synthesis kit as per manufacturer’s instructions (Thermo Scientific, Walter, MA, USA) (33).

### Gene expression analysis by qRT-PCR

The cDNA was diluted to 1:20 and analysed using iQ5 Real time PCR detection system (BioRad, Hercules, California, USA) with SYBR Green detection system. Each sample was set up in triplicates in a 96-well plate (BioRad, Hercules, California, USA) in a final reaction mixture of 10μl containing 2X SYBR iQ SYBR Green supermix and 10μM primer mix, along with the diluted cDNA template. The cycling conditions for the PCR were as follows: 95°C for 5mins, followed by 39 cycles of 95°C for 30s, 57°C for 30s and 72°C for 30s. The amplification specificity and the primer dimer were calculated by the melt curve acquired after 81 cycles of heating the PCR products from 55°C to 95°C for 20s, with a 0.5°C increase per cycle. The threshold cycles (Ct) were obtained from the iQ5 optical system software, and the relative quantities of transcripts were determined using the standard curve method with normalization performed against the mean of the reference control gene, *gmk.* The untreated WT cells, at the respective time points was normalized to 1, and all other samples were calculated as fold change to this reference value, using the 2^-^_ΔΔCt_ method (77).

### Atomic force microscopy

The pre-inoculum was grown overnight at 37°C. The absorbance of the bacterial pre-inoculum was normalised to 2.0 at 600 nm and 0.2% culture was added in 50 ml flasks and the cultures were incubated at 37°C at 160 rpm for 12 hours. Aliquots were obtained at 3 hours, 6 hours and 12 hours. The cells were pelleted down by centrifugation at 6000 rpm and washed with double distilled water thrice. Around 5 μl was placed on top of a glass coverslip and the images were obtained by using NX-10 AFM (Park systems, South Korea) (78, 79). Cell width was measured using the image processing software XEI (Park Systems, South Korea). For the Δ*rffG*Δ*rfbB* strain, which although are spherical but not perfect spheres, the cell width was measured considering the smaller dimension to be the width of the cell.

### Motility assays

Bacterial motility assays were performed as described previously (42). Briefly, swimming motility assay was performed on a freshly prepared 0.3% agar. Briefly, swimming motility assay was performed on a freshly prepared 0.3% agar, where 3μl of overnight grown cells whose OD at 600 nm was normalised to an absorbance of 2.0 were inoculated with a gentle stab at the middle of the plate. For swarming motility assays, 0.5% agar was used along with 0.5% glucose (HiMedia, Mumbai, India). Around 5μl of culture was placed on the surface and the middle of the plate for swarming motility assay. For both types of motility assays the plates were incubated at 37°C for 8 hours. The plates were imaged using the ImageQuant LAS4000 (GE Healthcare). The extent of area covered or spanned by the motile cells after 8 hours was quantified by the ImageJ software (1.53t).

### Adhesion, invasion and intracellular replication assay

HeLa cells were cultured in DMEM containing 4.5g/L D-glucose, 4mM L-glutamine and 1.5g/L sodium bicarbonate (HiMedia, Mumbai, India) supplemented with 10% (v/v) FBS, 5% CO_2_ at 37°C. Approximately 24 hours before initiating the experiment the cells were seeded at the density of 1.5×10^4^ cells/well in a 96-well plate. Bacterial cells were grown overnight at 37°C. The absorbance of the overnight grown pre-inoculum was normalised to OD 2.0 at 600 nm and 50μl of this pre-inoculum was inoculated in 50ml of LB and grown at 37°C for 10 hours (80, 81). For adhesion assay, both the inoculum and the 96-well plates were kept on ice for 15 minutes prior to infection. Subsequently, cells were infected at an MOI 1:10 and incubated on ice for 30 minutes. Cells were then washed thrice with sterile PBS, lysed using 100μl of 0.1% Triton X-100, and appropriate dilutions were plated on LB agar plates. For invasion and intracellular replication assays, HeLa cells were infected with an MOI 1:10 and 96-well plates were centrifuged at 2000 rpm of 2 minutes. Following incubation at 37°C for 50 minutes, the cells were washed twice with PBS and 100μl of 100μg/ml of gentamycin made in DMEM was added to kill the extracellular bacteria. To study invasion, cells were lysed with 0.1% Triton X-100 at 2 hours post infection, and appropriate dilutions were plated on LB agar. Following 2 hours, the gentamycin concentration was reduced to 25μg/ml and cells were incubated at 37°C. At 18 hours post infection, gentamycin-containing medium was aspirated and the cells washed twice with PBS followed by the addition of 0.1% Triton X-100 to the wells. Appropriate dilutions were plated on LB agar plates to determine the intracellular bacterial load.

### Mice infections

Six-to-eight week old, male, C57BL/6 mice were orally infected with 1×10^8^ CFU/mouse of either the *S*. Typhimurium WT or the Δ*rffG*Δr*fbB* strain (82). Organ bacterial burden was estimated 4 days post infection. The organs were harvested, weighed, and homogenized in 1 ml sterile PBS. Appropriate dilutions were plated on LB agar plates to enumerate CFU. Data were represented as log_10_(CFU/gram tissue weight).

### Measurement of cytokines in sera

The amounts of TNFα, IL6 and IFNγ in sera were quantified using sandwich ELISA (83) according to manufacturer’s instructions using the respective cytokine estimation kits (ThermoFisher Scientific, USA). 3,3’,5,5’-Tetramethylbenzidine (TMB) was used as the chromogenic substrate, and absorbance was recorded at 450 nm using VersaMax Microplate Reader (Molecular Devices, USA). Appropriate standards (31.25-2000pg/ml) provided with the respective cytokine kits were used to estimate the standard absorbance values (at 450nm) to obtain the absorbance versus concentration dose-response curves. The cytokine amounts (unknown/test) from sera samples were interpolated from the respective standard curves within the detection range.

### Statistical analysis

All graphs were plotted, and the statistical analyses performed using GraphPad Prism 8 (v 8.0.2) software (GraphPad, La Jolla, CA). For most of the experiments, statistical analyses were performed using either one-way or two-way ANOVA. Data are represented as mean ± SEM, where * *p* < 0.05, ∗∗ *p* < 0.01, ∗∗∗ *p* < 0.001 and ∗∗∗∗ *p* < 0.0001. For *in vivo* infection studies, statistical analyses of mice survival curves post infection were performed using the log-rank (Mantel-Cox) test. For estimation of cytokines, statistical analysis was performed using the unpaired student *t* test, where * *p* < 0.05; ** *p* < 0.01 and *** *p* < 0.001.

## Acknowledgements

Imaging studies were performed at the Bioengineering Imaging and AFMM facilities in IISc. We thank the help of the Central Animal Facility, IISc for all the experiments involving mice. Finally, the support from all members of the DpN lab is greatly appreciated.

## Conflict of interest

The authors declare that the research was conducted without any commercial or financial relationships that could be construed as a potential conflict of interest.

## Funding information

This work was supported by core grants from IISc and the DBT-IISc partnership program. In addition, the infrastructural support from DST-FIST and UGC CAS/SAP is greatly appreciated.

## Notes

### Competing Interest Statement

The authors have declared no competing interest.

