## Supplemental Information for "Functional loss of *rffG* and *rfbB,* encoding dTDP-glucose 4,6-dehydratase, changes colony morphology, cell shape, motility and virulence in *Salmonella* Typhimurium"

**Supplementary Information**

| Condition | *rffG* in FC in D23580 | *rfbB* FC in D23580 | *hilA* in D23580 | *hilD* in D23580 | *ssaG* in D23580 |
|---|---|---|---|---|---|
| **EEP** | 1 | 1 | 1 | 1 | 1 |
| **MEP** | 0.88 | 0.71 | 0.95 | 0.79 | 1 |
| **LEP** | 0.78 | 0.42 | **16.41** | **12.57** | 1 |
| **ESP** | 0.64 | 0.28 | **18.64** | **11.38** | **13.34** |
| **LSP** | 0.17 | 0.04 | 0.95 | 1.01 | 1.79 |
| **MEP** | 1 | 1 | 1 | 1 | 1 |
| **NaCl shock** | 0.3 | 0.25 | 1 | 2.73 | 1 |
| **Bile shock** | 1.63 | 1.32 | 1 | 0.9 | 1 |
| **Low $Fe^{2+}$ shock** | 0.92 | 0.37 | 1 | 1.13 | 1 |
| **Anaerobic shock** | 1 | 0.18 | 1.06 | **4.67** | 1 |
| **Anaerobic growth** | 1 | 1 | 1 | 1 | 1 |
| **Oxygen shock** | 1.84 | 1.45 | **3.92** | 1.56 | 1 |
| **InSPI2** | 1 | 1 | 1 | 1 | 1 |
| **Peroxide shock(InSPI2)** | 0.26 | 0.17 | 1 | 0.32 | 0.57 |
| **Nitric Oxide shock(InSPI2)** | 0.78 | 0.43 | 1 | 0.62 | 0.89 |
| **NonSPI2** | 1 | 1 | 1 | 1 | 1 |
| **InSPI2** | 0.79 | 0.6 | 0.72 | 0.97 | **136.47** |
| **ESP** | 1 | 1 | 1 | 1 | 1 |
| **Macrophage** | 1.55 | 1.2 | 0.05 | 0.2 | **10.07** |

**Table S1: Intra-strain relative expression levels of *rffG* and *rfbB* genes in *S.* Typhimurium D23580 strain (SalCom V2.0).** Tabular representation of the intra-strain expression levels of *rffG* and *rfbB* under a suite of experimental conditions. The genes, *hilA* and *hilD* have been tabulated as reference genes from the SPI-1 island and *ssaG* as a reference gene from the SPI-2 island.

Abbreviations: EEP: Early exponential phase; MEP: Mid exponential phase; LEP: Late exponential phase; ESP: Early stationary phase; LSP: Late stationary phase; Non SPI2: PCN medium (pH 7.4, 25 mM $P_i$); InSPI2: PCN medium (pH 5.8, 0.4 mM $P_i$).

**Table S2: List of bacterial strains used in this study.**

| Sl. No. | Name of the strain | Source |
|---|---|---|
| 1. | *Salmonella* Typhimurium 14028s | (1) |
| 2. | *Salmonella* Typhimurium Δ*rffG* | This study |
| 3. | *Salmonella* Typhimurium Δ*rfbB* | This study |
| 4. | *Salmonella* Typhimurium Δ*rffG*Δ*rfbB* | This study |
| 5. | *Salmonella* Typhimurium WT/pACDH | This study |
| 6. | *Salmonella* Typhimurium WT/*rffG* | This study |
| 7. | *Salmonella* Typhimurium WT/*rfbB* | This study |
| 8. | *Salmonella* Typhimurium Δ*rffG*Δ*rfbB*/pACDH | This study |
| 9. | *Salmonella* Typhimurium Δ*rffG*Δ*rfbB/rffG* | This study |
| 10. | *Salmonella* Typhimurium Δ*rffG*Δ*rfbB/rfbB* | This study |

**Table S3: List of primers used in this study.**

| Oligonucleotides (5’ → 3’) | | |
|---|---|---|
| **Knockout generation primers** | | |
| ***rffG:: Kan$^r$*** | FP | AAGGAGTCTGGCGCTGATGAAACGCATTCTGGTGACCGGCGTGTAGGCTGGAGCTGCTT |
| | RP | TTAGCGTTTCAGTCCTAAGCGTTCGCCCTGATAACTGCCAGTCCATATGAATATCCTCCTTAG |
| ***rfbB:: Chl$^r$*** | FP | ATGGAATAGAAAAGTGAAGATACTTATTACTGGCGGGGCAATGTAGGCTGGAGCTGCTTCG |
| | RP | TTACTGGCGTCCTTCATAGTTCTGTTCTATCCAACTCTGACGGCTGACATGGGAATTAGC |
| **Knockout confirmation primers** | | |
| ***rffG*** | FP | GTCGAACCGAATATCCGTCAG |
| | RP | CCAGACAGGCAATTTTGAAGC |
| ***rfbB* (Chl$^r$ internal primers)** | FP | GTGGTATTCACTCCAGAGC |
| | RP | CCGTTGATATATCCCAATGGC |
| **Gene cloning primers** | | |
| ***rffG*** | FP | CATGCCATGGATGAAACGCATTCTGGTGACC |
| | RP | CCCAAGCTTTTAGCGTTTCAGTCCTAAGCG |
| ***rfbB*** | FP | CATGCCATGGGTGAAGATACTTATTACTGGCG |
| | RP | CCCAAGCTTTTACTGGCGTCCTTCATAGTTC |

| Oligonucleotides (5' → 3') | | |
|---|---|---|
| **Gene sequencing primers** | | |
| ***rffG*** | | |
| ***Fragment 1*** | FP | CACAGGAAACAGCTATGACC |
| | RP | GCGTAGGCAGACCGTAGGTA |
| ***Fragment 2*** | FP | TTCCGCTTCCACCATATCTC |
| | RP | GCGTAGGCAGACCGTAGGTA |
| ***Fragment 3*** | FP | TTCCGCTTCCACCATATCTC |
| | RP | GTGCCAACATAGTAAGCCAG |
| ***rfbB*** | | |
| ***Fragment 1*** | FP | CACAGGAAACAGCTATGACC |
| | RP | GATCCGCGACATAAGTG |
| ***Fragment 2*** | FP | GGTGATGCATTTGGC |
| | RP | GTGCCAACATAGTAAGCCAG |

| Oligonucleotides (5' → 3') | | |
|---|---|---|
| **qPCR primers** | | |
| ***hilA*** | FP | TAATCGTCCGGTCGTAGTGG |
| | RP | TGCGGCAGTTCTTCGTAATG |
| ***hilD*** | FP | AACGTGACGCTTGAAGAGGT |
| | RP | GAACGCCGTTTTCAGATGTT |
| ***sipC*** | FP | CGCGAATACGTTAATGCTGA |
| | RP | CGCGCTCTGGGAAATACTAC |
| ***flhD*** | FP | ATCGTCCAGGACAAAGCATC |
| | RP | TCGTCCACTTCATTGAGCAG |
| ***fliC*** | FP | TGACAGCAGCAGGTGTTACC |
| | RP | CGCCACCCAGTTTGTTTAGT |
| ***fljB*** | FP | GCCAACGACGGTGAAACTAT |
| | RP | CACCCGTAGCCGCTTTAATA |
| ***gmk*** | FP | TTCCGTTTCACATACCACGC |
| | RP | CCTGCCAGTCGATATCCAGA |

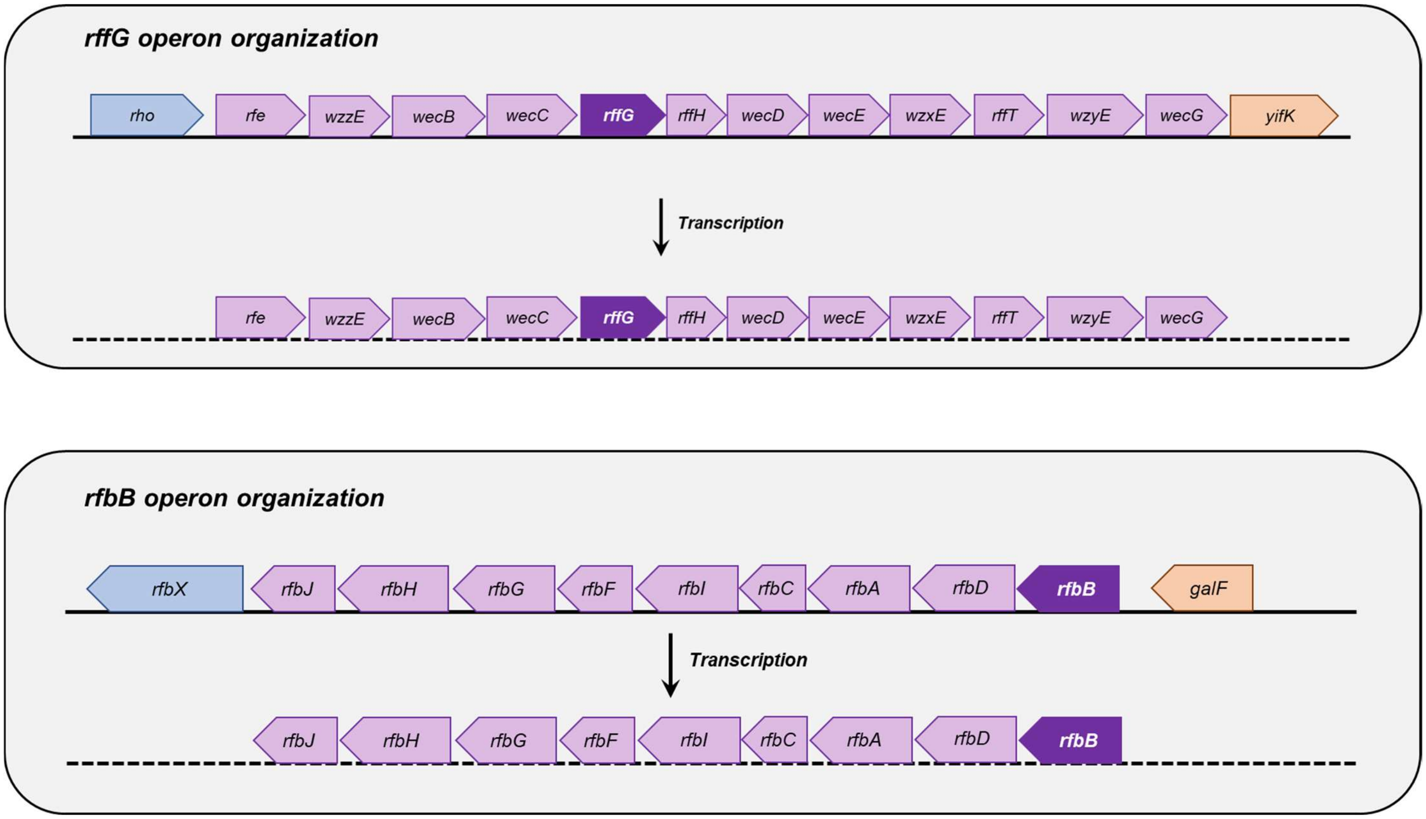


**Figure S1. Operon organization of *rffG* and *rfbB* in *S.* Typhimurium LT2 genome.** The genomic location and organization of the genes, *rffG* and *rfbB* was explored in the BioCyc database. *Salmonella* Typhimurium LT2 was selected as the query organism for this analysis. BioCyc.org is a microbial genome web portal that combines several genomes with additional information inferred by computer programs, imported from other databases and curated from biomedical literature by biologist curators. BioCyc also provides an extensive range of query tools, visualization services and analysis software (2).

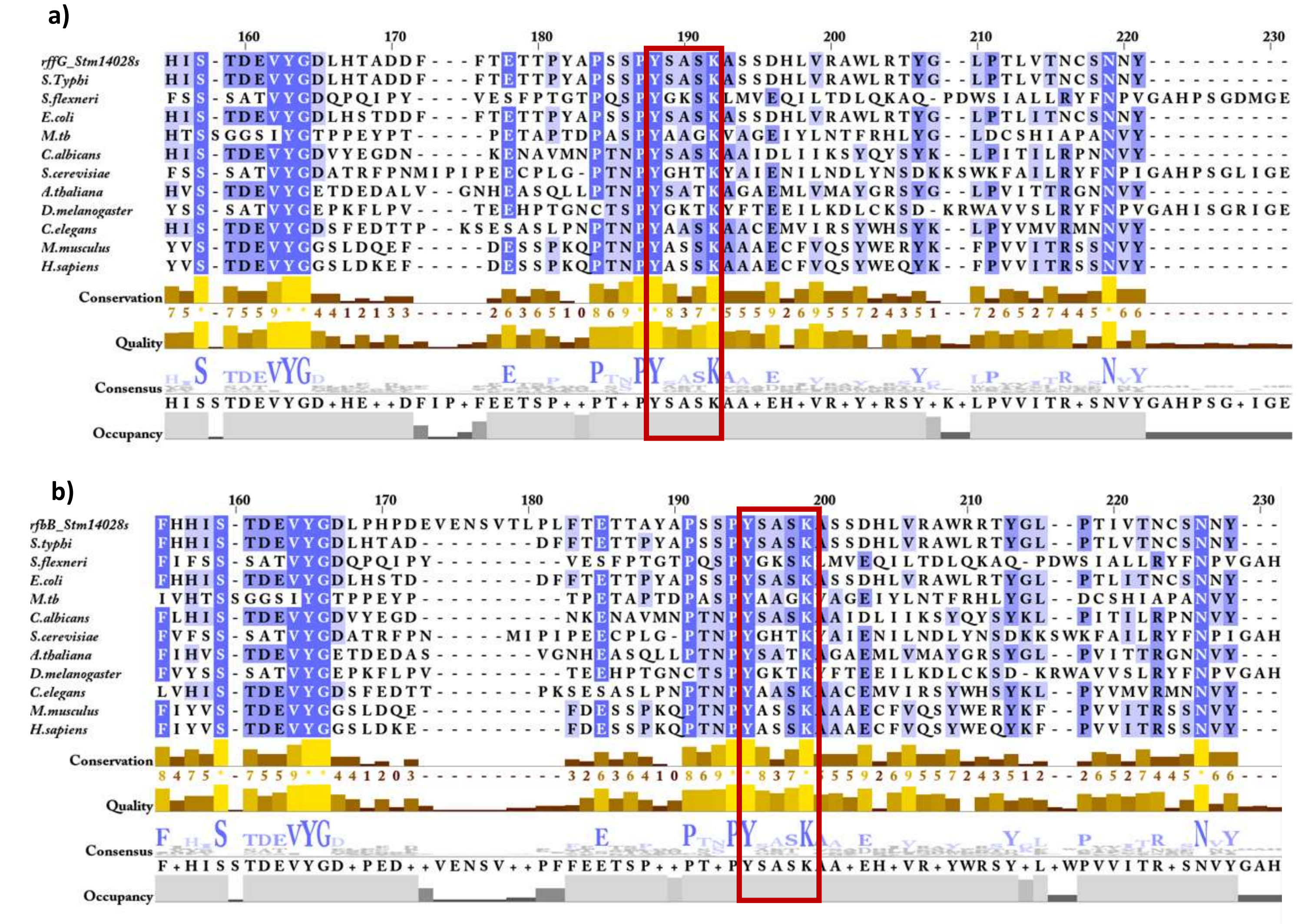


**Figure S2. Multiple sequence alignment (MSA) of homologs with highest percentage identity to *S.* Typhimurium 14028s encoded proteins, RffG and RfbB.** A MSA was performed using CLUSTAL Omega program and represented in JalView with homologs of **(a)** RffG and **(b)** RfbB proteins encoding the enzyme, dTDP glucose 4,6-dehydratase in *S.* Typhimurium 14028s. The reference genomes from these representative organisms have been used for performing this analysis: *Salmonella enterica subsp. enterica serovar* Typhi str. CT18, *Shigella flexneri*, *Escherichia coli*, *Mycobacterium tuberculosis*, *Candida albicans, Saccharomyces cerevisiae*, *Arabidopsis thaliana, Drosophila melanogaster, Caenorhabditis elegans, Mus musculus and Homo sapiens.* The conserved catalytic motif YXXXK in these proteins have been highlighted in the red box.

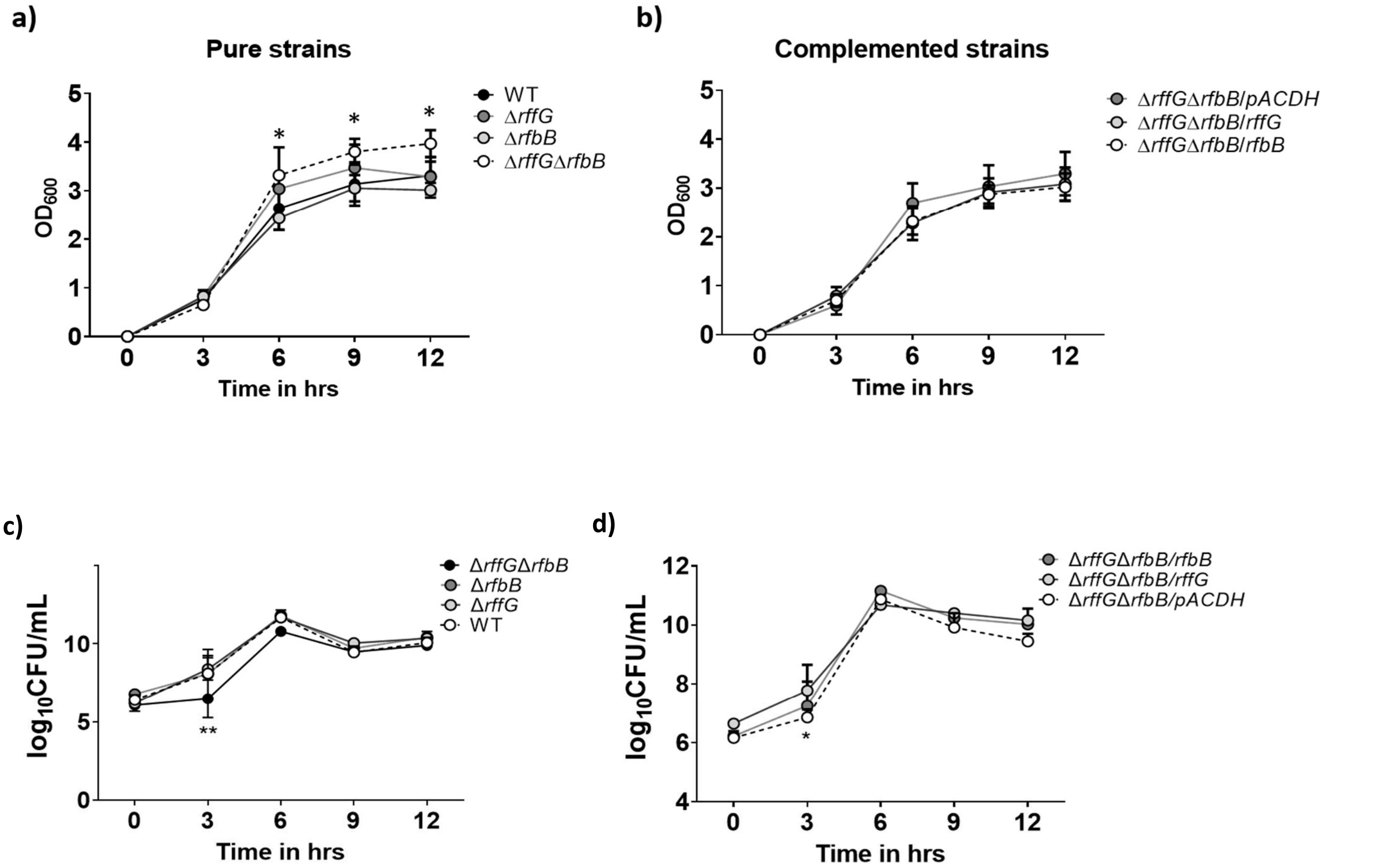


**Figure S3. *S.* Typhimurium Δ*rffG*Δ*rfbB* shows higher O.D. as compared to the WT and the single gene-deleted mutants. (a)** *S.* Typhimurium WT, Δ*rffG*, Δ*rfbB* and Δ*rffG*Δ*rfbB* were grown in LB broth for the indicated time points. **(b)** Growth of *S.* Typhimurium Δ*rffG*Δ*rfbB* with plasmid alone or either *rffG* or *rfbB* expressed *in trans*. Briefly, overnight grown cultures were normalized to O.D. 2 at 600 nm and 0.2% bacterial culture was inoculated in an Erlenmeyer flask containing 50 ml LB broth. The flasks were incubated at 37ºC under shaking conditions (160 rpm) for 12 hours. At indicated time intervals, 1 ml of the bacterial culture from each flask was aspirated and the O.D. was measured at 600 nm. **(c)** *S.* Typhimurium WT, Δ*rffG*, Δ*rfbB*, Δ*rffG*Δ*rfbB* and **(d)** complemented strains, Δ*rffG*Δ*rfbB*/pACDH, Δ*rffG*Δ*rfbB/rffG*, Δ*rffG*Δ*rfbB/rfbB* were grown in 50 ml LB broth at 37ºC. Culture aliquots were aspirated at indicated time points and appropriate dilutions were plated on LB agar plates. The plates were incubated at 37ºC, and the colonies obtained were enumerated. Data shown are representative of 3 independent experiments, plotted as mean ± SEM. Statistical analysis was performed using two-way ANOVA, where * $p < 0.05$ and ** $p < 0.01$.

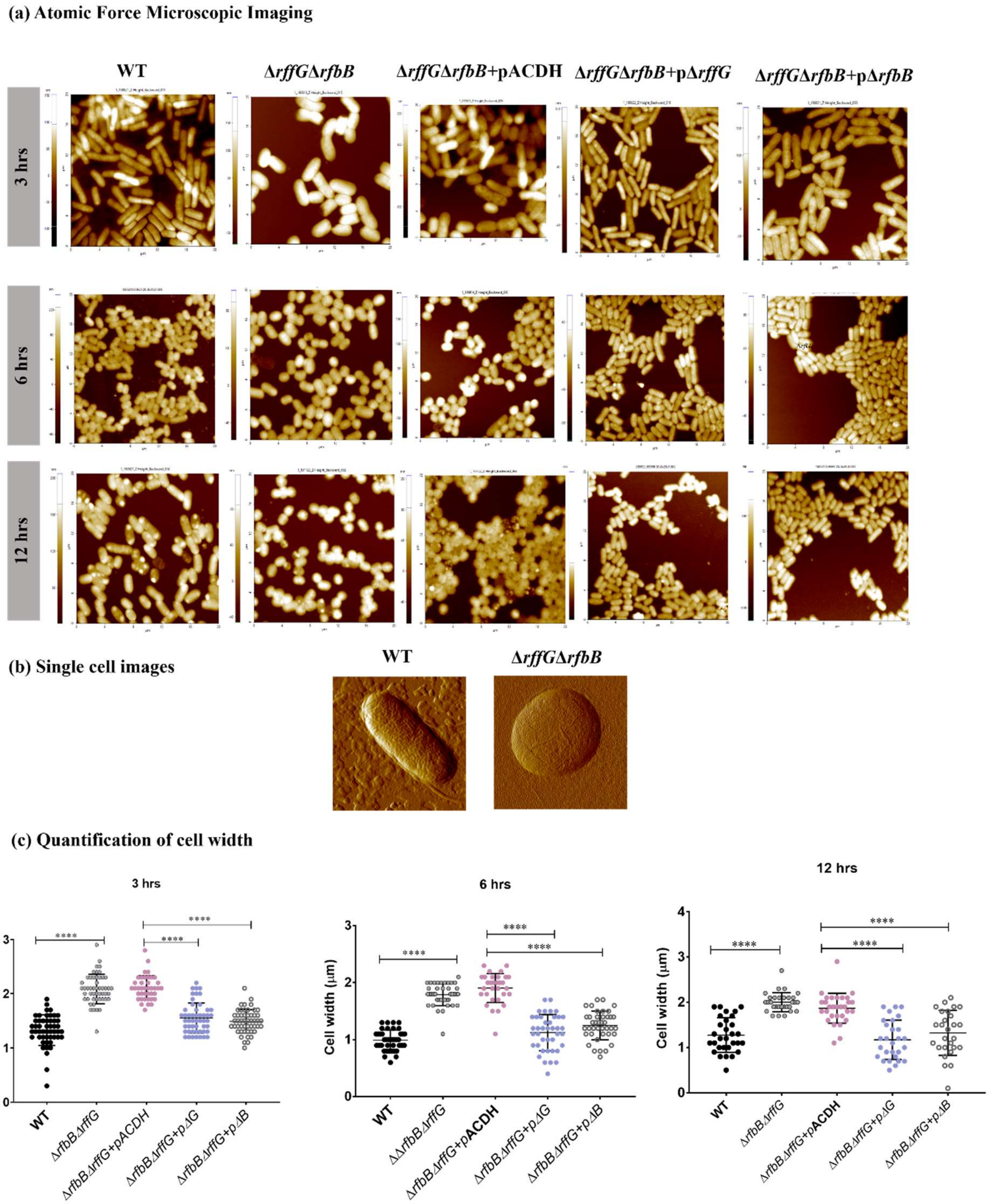


**Figure S4. Complementation of *S*. Typhimurium Δ*rffG*Δ*rfbB* strain with WT copy of *rffG* or *rfbB* restores cell width. (a)** The bacterial strains were grown for the indicated time points (3, 6 or 12 hours) and AFM images were acquired in the non-contact mode. **(b)** Single cell images of the WT and the Δ*rffG*Δ*rfbB* at 6 hours. **(c)** Quantification of the cell width was performed and for each condition, the width of at least 50 cells was determined. Data are representative of 3 independent experiments plotted as mean ± SEM. Statistical analysis was performed using two-way ANOVA, where * $p < 0.05$; ** $p < 0.01$; *** $p < 0.001$ and **** $p < 0.0001$.

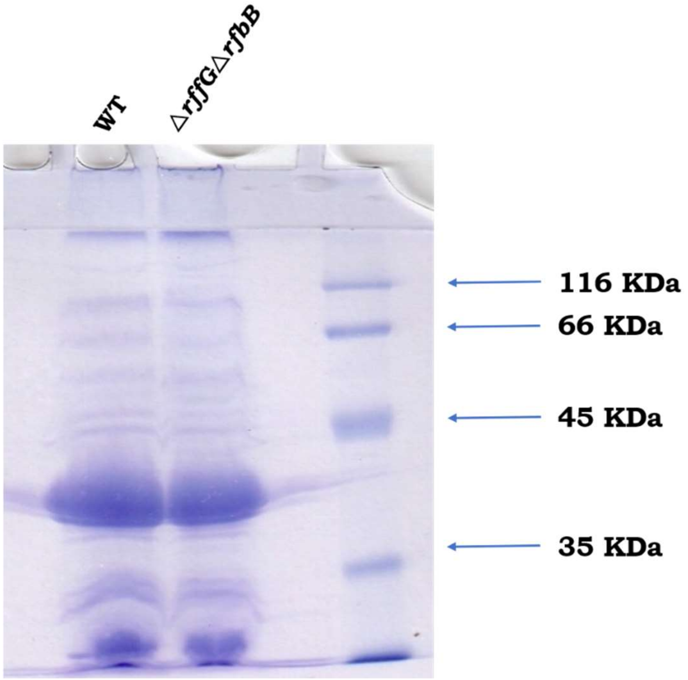


**Figure S5. Outer membrane protein (OMP) profile of *S*. Typhimurium WT and Δ*rffG*Δ*rfbB*.** *S*. Typhimurium WT and Δ*rffG*Δ*rfbB* were grown at 37ºC for 15 hours. The OMPs were isolated and resolved on a 12% SDS PAGE.

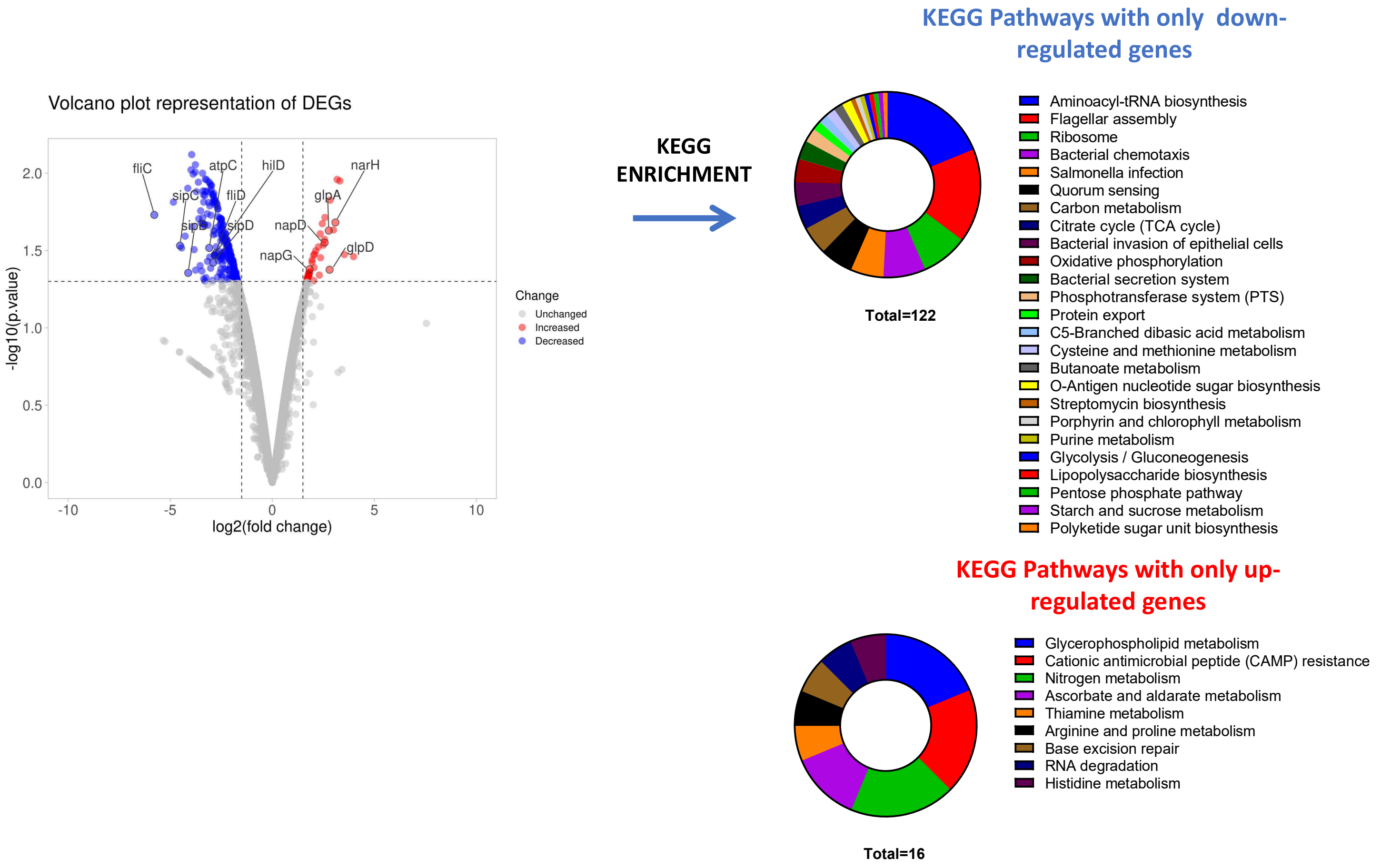


**Figure S6. RNAseq analysis depicts significantly more downregulated pathways in the *S.* Typhimurium Δ*rffG*Δ*rfbB* strain. (a)** Volcano plot representation of Differentially Expressed Genes ($-1.5 \leq \log_2$Fold Change $\geq 1.5$) in the Δ*rffG*Δ*rfbB* strain. **(b)** KEGG pathway enrichment of the significantly upregulated or downregulated genes in the Δ*rffG*Δ*rfbB* strain of *S*. Typhimurium.

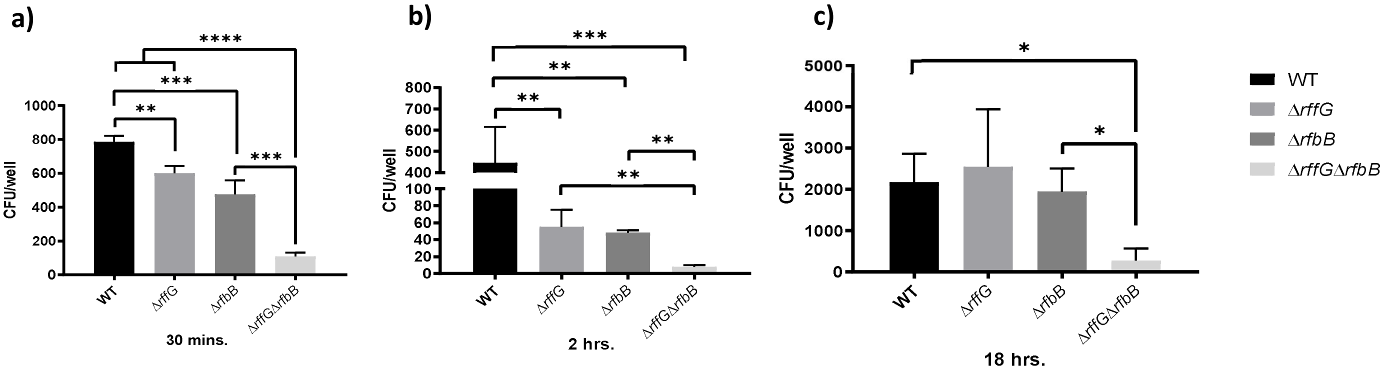


**Figure S7. *S.* Typhimurium Δ*rffG,* Δ*rfbB* and Δ*rffG*Δ*rfbB* are compromised with respect to adhesion and invasion in HeLa cells**. HeLa cells were infected at an MOI 1:10 and **(a)** adhesion, **(b)** invasion and **(c)** intracellular replication were studied at 30 minutes, 2 hours, and 18 hours post infection, respectively. Data are representative of 3 independent experiments expressed as mean ± SEM. Statistical analysis was performed using two-way ANOVA, where * $p < 0.05$; ** $p < 0.01$; *** $p < 0.001$ and **** $p < 0.0001$.

### SUPPLEMENTARY METHODS

**Multiple Sequence Alignment:** The protein sequences, RfbB (361aa; locus id: STM14_2591) and RffG (355aa; locus id: STM14_4720) from *S*. Typhimurium 14028s were obtained from the NCBI Protein records. These sequences were independently used as query sequences to perform protein BLAST (using the NCBI BLASTp suite) against the non-redundant (NR) sequence database. To identify close homologs, the search set was curated and limited to records from reference genomes of prokaryotic and eukaryotic representative organisms. BLAST was performed with the default algorithm parameters. The hits obtained from the above analysis were further filtered and only homologs with highest percentage identity in each reference genome were taken for further analysis. These FASTA sequences producing significant alignment with the input query sequence were downloaded and used as an input in the CLUSTAL Omega program (3). CLUSTAL Omega is an MSA program that uses seeded guide trees and HMM profile-profile techniques to generate alignments between three or more sequences. For visualization and analysis of the alignment JalView (4) was used to highlight important features in the alignment, e.g., the conserved residues, and generating a consensus sequence.

**Library preparation, RNA sequencing, data analysis and KEGG pathway enrichment:** For each sample, 1µg of total RNA was taken as input for RiboMinus Transcriptome Isolation Kit (Thermo Fisher Scientific) to remove ribosomal RNA. The ribosomal RNA depleted samples were used to generate a sequencing library using NEB NEXT RNA Ultra II library preparation kit for Illumina. Briefly, RNA was fragmented, and reverse transcribed to generate cDNA. Hairpin adapters were ligated to fragmented double-strand cDNA and USER enzyme was used to cleave the hairpin structure. Ampure beads were used to purify adapter-ligated fragments and the purified product was amplified using Illumina Multiplex Adapter primers to generate a sequencing library with barcodes for each sample. The library was quantitated using Qubit DNA High Sensitivity quantitation assay and library quality was checked on the Bioanalyzer 2100 using the Agilent 1500 DNA Kit. The QC passed libraries were diluted to 2nM and pooled together. The pooled library was further diluted to sequence it on Illumina Hiseq as per the manufacturer's recommendation. The Hiseq Control Software was used to setup 2x150 bp run on Hiseq and data was demultiplexed using bcl2fastq v2.1.9. The sequence data quality was checked using FastQC and MultiQC (5) software. The data was checked for base call quality distribution, % bases above Q20, Q30, %GC, and sequencing adapter contamination. All samples passed the QC threshold (Q30 > 80%). Raw sequence reads were processed to remove the adapter sequences and low-quality bases using Trim Galore. Alignment and expression analysis: The QC passed reads were mapped onto indexed *S*. Typhimurium 14028s strain reference genome using HISAT2 (6, 7) aligner. On average 97.53% of the reads aligned onto the reference genome. The PCR and optical duplicates were marked and removed using Picard tools (Broad Institute). Gene level expression values were obtained as read counts using featureCounts software (8). For differential expression analysis, DESeq2 (9, 10) package was used. Genes with < 5 reads in any one of the samples per condition were removed. The read counts were normalized (variance stabilized normalized counts) and differential expression analysis was performed. All the samples were compared to WT untreated condition independently. Genes with absolute log2 fold change ≥ 1.5 and p-value ≤ 0.05 (Wald's test) were considered significant. The expression profile of the DEGs across the samples are presented as volcano plots using the VolcaNoseR web app (11). Gene set enrichment analysis was performed with the significant DEGs for pathway enrichment using the KEGG mapper tool (12, 13).

**Outer membrane protein isolation, purification, and quantitation:** OMPs were isolated from the *S.* Typhimurium strains grown in LB (14). Briefly, cells were harvested in the late-log phase (12 hours) of growth and washed twice with 1X PBS. Approximately O.D. 4 (600 nm) cells were used for the extraction. OMP concentrations were determined by the Bradford's assay, using BSA as standard. Equal volume of resuspended solution was loaded and analyzed by a 12.5% SDS-PAGE and visualized by staining with Coomassie Brilliant Blue (Sigma).
